# MondrianMap: Navigating Gene Set Hierarchies with Multi-Resolution Enrichment Maps

**DOI:** 10.64898/2026.06.27.735020

**Authors:** Fuad Al Abir, Zongliang Yue, Ehsan Saghapour, Md Delower Hossain, Zhandos Sembay, Jake Y. Chen

## Abstract

Gene Ontology encodes genes as a hierarchy, yet every enrichment visualization flattens it into a ranked list, discarding the ability to view the same process at different levels of abstraction. We present MondrianMap, a free interactive web application (https://mondrianmap.smartdrugdiscovery.org/) that organizes enrichment results into 13 semantically principled layers derived from the GOALS framework and renders them as color-encoded rectangular maps where block area reflects significance, color encodes effect direction, and spatial proximity preserves semantic relations, all within an interactive interface. Three case studies across the NIH Common Fund Data Ecosystem demonstrate that visualization facilitates recognition of patterns that are difficult to discern in flat outputs: (1) LINCS CRISPR perturbations reveal that TP53 and KRAS knockouts produce opposite color maps at a single semantic layer, the same immune recruitment processes suppressed by TP53 loss are activated by KRAS disruption; (2) GTEx aging signatures expose the inflammaging paradox as an immediate visual phenomenon, identical antimicrobial defense programs appear uniformly upregulated in aging blood yet uniformly downregulated in aging liver at matched semantic resolution; and (3) MoTrPAC exercise data capture temporal dynamics as color transitions where brown adipose tissue undergoes a threshold switch from two enriched terms to thirty-four at a single molecular layer, while cardiac tissue reverses from uniformly activated to uniformly suppressed glycolytic metabolism as adaptation progresses. MondrianMap facilitates hierarchical visual reasoning, complementing statistical enrichment reporting for biological discovery.

**HIGHLIGHTS:**

- MondrianMap provides multi-resolution enrichment visualization for Gene Ontology
- Layer-specific views reveal directional oppositions difficult to discern in flat term lists
- Navigating layers organizes one enrichment into a multi-scale biological narrative
- Demonstrated on cancer, aging, and exercise data across three CFDE programs

**IN BRIEF:** MondrianMap is a web application that transforms gene set enrichment results into layered visualizations encoding regulation, significance, and semantic hierarchy. Across cancer, aging, and exercise datasets, viewing enrichment at defined semantic layers exposes directional inversions, temporal switches, and multi-scale biological narratives that are difficult to extract from conventional flat enrichment outputs.

**THE BIGGER PICTURE:** When researchers measure gene expression changes in disease, aging, or drug response, they rely on enrichment analysis to translate thousands of molecular measurements into interpretable biological themes. The standard output is a ranked list of processes sorted by statistical significance. This format served the field well when studies examined a single condition; however, modern genomics routinely compares dozens of tissues, timepoints, and perturbations, each generating hundreds of enriched terms. The critical limitation is not statistical power; however, interpretive structure: a flat list cannot show whether two conditions activate the same biological process in opposite directions, whether a process visible at one level of abstraction disappears or transforms at another, or how a tissue’s functional response evolves across time.

These are precisely the questions that define contemporary systems biology. Does a tumor suppressor gene silence the same immune program that an oncogene activates? Does aging drive the same defense pathway upward in the blood and downward in the liver? Does an exercise response flip from activation to suppression as the tissue adapts? Answering these questions requires a visualization framework that preserves hierarchy, encodes direction, and enables comparison at matched levels of biological resolution. MondrianMap provides this framework by organizing Gene Ontology terms into quantitatively defined semantic layers and rendering enrichment as color-encoded rectangular maps navigable from molecular mechanism to system-level theme. The result is a tool that extends enrichment analysis from a reporting step into an interactive framework for biological reasoning, hypothesis generation, and cross-dataset discovery.

## INTRODUCTION

Gene set enrichment analysis is the standard method for translating differential expression into biological understanding. Tools such as Enrichr [1,2], g:Profiler [3], and DAVID [4] map gene lists onto annotated biological processes and rank the results by statistical significance, leaving the investigator to scan for coherent themes. This workflow has powered thousands of studies, yet it harbors a structural limitation that grows more consequential as datasets scale: the Gene Ontology is a directed acyclic graph, a multi-level hierarchy of biological abstraction; however, most conventional visualizations present it in a *flat* view. Dot plots, bar charts, ranked tables, and semantic similarity networks all collapse the ontology’s depth, and with that collapse, a critical capability is lost: the ability to view the same enrichment at a defined level of biological specificity, compare it across conditions at that level, and then move to a different level to see how the interpretation transforms.

Existing tools have addressed components of this problem. REVIGO [12] reduces redundancy through semantic similarity clustering, projecting terms onto a two-dimensional plane, elegant for small enrichment sets, however, the 2D projection collapses hierarchical depth, placing a specific enzymatic mechanism and a broad umbrella process in the same plot region if they share semantic neighbors. Enrichment Map [13] constructs gene-overlap networks in Cytoscape that reveal crosstalk structure, however, at the scale of modern studies, hundreds of terms per contrast, the network becomes a dense tangle that obscures rather than clarifies. Treemap and sunburst viewers faithfully represent the GO graph however, cannot simultaneously encode enrichment statistics, effect direction, and cross-condition comparison without visual overload. clusterProfiler [14], a widely used R package, offers semantic similarity grouping and ridge plots for multi-contrast comparison however, relies on data-driven clusters rather than biologically principled strata, and cannot show the investigator what happens to the same enrichment when viewed at a different semantic resolution. None of these tools was built to solve the joint problem of redundancy, hierarchy, effect direction, and multi-resolution comparison.

A principled solution requires a rigorous decomposition of the Gene Ontology into semantic layers, strata with consistent biological meaning at each level, across the entire ontology. The GOALS (Gene Ontology Analysis with Layered Shells) framework [15] provides this foundation. GOALS applies an Initiation-Assignment-Expansion-Repetition (IAER) algorithm to GO co-membership networks, assigning each GO-BP term to one of 13 semantic layers (L1–L13) such that the distrihowever,ion achieves a power-law fit with R² = 0.98, reducing mean sample variance of gene set sizes by two orders of magnitude compared to conventional depth-based stratification. Layers 1–3 contain mechanistically precise terms, specific signaling cascades, enzymatic reactions, vesicle fusion events, while layers 11–13 capture umbrella themes such as regulation of cell proliferation and inflammatory response. This stratification transforms the ontology from a monolithic graph into a navigable hierarchy with quantitatively defined resolutions.

Building on GOALS, we developed MondrianMap, a free, interactive web application (https://mondrianmap.smartdrugdiscovery.org/) that transforms enrichment outputs into a hierarchically navigable visualization. The concept of using nested rectangles to encode multiple dimensions of biological data was inspired by Piet Mondrian’s abstract compositions of colored rectangles. MondrianMap integrates five components: (i) GOALS-based 13-layer decomposition for principled multi-resolution navigation from umbrella theme to mechanistic detail; (ii) GoBERT-based semantic embeddings [30] of GO terms combined with UMAP dimensionality reduction [31] to position related terms in spatial proximity while preserving semantic structure; (iii) dual color-encoding of enrichment direction (red for upregulated, blue for downregulated genes) alongside effect size (block area proportional to −log₁₀(adjusted p-value)), treating directionality as a first-class visual variable rather than a supplementary annotation; (iv) Jaccard-based crosstalk edges connecting GO terms that share genes, revealing functional modules and candidate hub genes; and (v) an integrated AI hypothesis generation module that synthesizes layer-specific biological narratives grounded in the GOALS context of each layer.

We demonstrate MondrianMap’s utility across three case studies drawn from NIH Common Fund Data Ecosystem (CFDE) programs: LINCS L1000 CRISPR perturbations (cancer pharmacogenomics) [6,7], GTEx aging signatures (tissue-specific aging biology) [8,9], and MoTrPAC endurance training (exercise temporal dynamics) [10,11]. In each case, layered MondrianMaps facilitate recognition of patterns that are difficult to discern in flat enrichment outputs: directional inversions between cancer drivers at a single semantic layer, the inflammaging paradox rendered as a color contrast across tissues, temporal threshold switches and directional reversals in exercise adaptation, and multi-resolution narratives that transform coherently as one traverses the GOALS hierarchy. These findings demonstrate that the joint encoding of hierarchy, direction, and semantic proximity within a single interactive framework provides a visual reasoning framework that complements statistical enrichment reporting for biological discovery.

## RESULTS

### MondrianMap Architecture and Features

MondrianMap is an interactive web application that translates gene sets into enriched and hierarchical visualizations. The pipeline transforms raw gene sets, whether provided via custom user queries or preloaded CFDE datasets, into mechanistic insight through four functional modules: a. data ingestion and GO enrichment, b. GOALS layer assignment, c. semantic layout, and d. AI-powered hypothesis generation. At its core, MondrianMap performs GO Biological Process enrichment via Fisher’s exact test with Benjamini–Hochberg FDR correction (p ≤ 0.05), assigns enriched terms to the GOALS framework spanning 13 biological resolution levels (L1–L13), and positions terms using semantic embeddings (GoBERT) projected into two-dimensional space via UMAP, clustering functionally related processes while reducing visual redundancy. Block area encodes statistical magnitude (−log□□(adjusted p-value)), while directional color (red for upregulated, blue for downregulated, yellow for bidirectional/shared) immediately conveys the sign of regulation. Crosstalk edges link GO term pairs sharing genes at a Jaccard index ≥ 0.15, creating a visual graph of biological mechanisms interconnected within each layer and across the hierarchy. We presented the overall pipeline of the MondrianMap application in Figure 1.

**Figure 1.**
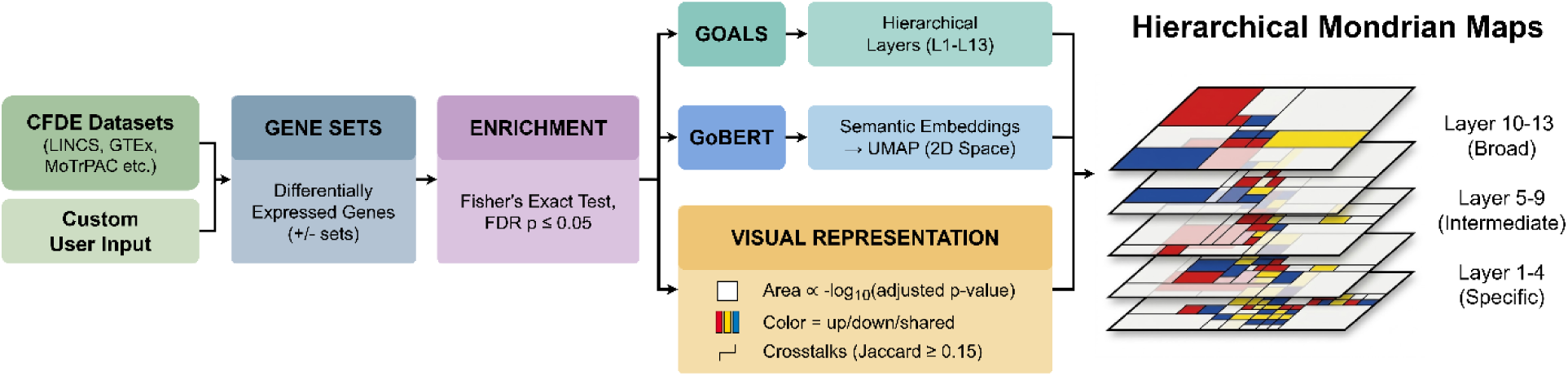
Overview of the MondrianMap workflow. Differentially expressed gene sets from user query or CFDE datasets (LINCS, GTEx, MoTrPAC) are processed through enrichment analysis (Fisher’s exact test), producing gene set statistics for GO Biological Process terms. GOALS layer assignment organizes enriched terms into 13 semantic layers: Layers 1–4 (Specific) capture molecular mechanisms (signaling cascades, protein modification); Layers 5–9 (Intermediate) encompass cellular processes (cell death, differentiation); Layers 10–13 (Broad) represent system-level themes (metabolism, development, reproduction). GoBERT embeddings initialize semantic proximity for spatial layout in a 1024-dimensional space, and UMAP generates 2D coordinates for each term, preserving the semantic distance. Hierarchical decomposition creates multi-resolution MondrianMaps: areas encode term significance, and color encodes effect direction (red = upregulated, blue = downregulated), providing clarity and insight, augmented by AI generated hypothesis module. Zoomable interface allows efficient navigation across all hierarchical layers.

The user interface facilitates a natural exploratory workflow (Figure 2) by offering two distinct input methods to trigger gene set enrichment analysis. Users can either input custom differentially expressed gene sets (Figure 2a, Custom Input) or select preloaded CFDE datasets and experiments (Figure 2a, Case Study). The main canvas (Figure 2b) displays the generated MondrianMap visualization, with any individual GOALS layer or all the 13 layers combined, at the user’s selection. The rightmost panel shows the enrichment results (Figure 2c) with all significant terms, p-values, gene counts, and layer assignments. Filter controls allow real-time refinement: by gene/crosstalk count (Figure 2d) to isolate high-confidence core processes, or by p-value/Jaccard threshold (Figure 2e) to trade specificity for breadth. Upon selecting one or more GO terms on the MondrianMap canvas or the enrichment results tables, an AI hypothesis can be generated for the selection, synthesizing a structured narrative (Figure 2f–g). All visualizations can be exported as PNG/SVG, enrichment results as JSON, and AI hypotheses with the selected term visualization as markdown. The application is freely accessible at https://mondrianmap.smartdrugdiscovery.org/.

**Figure 2.**
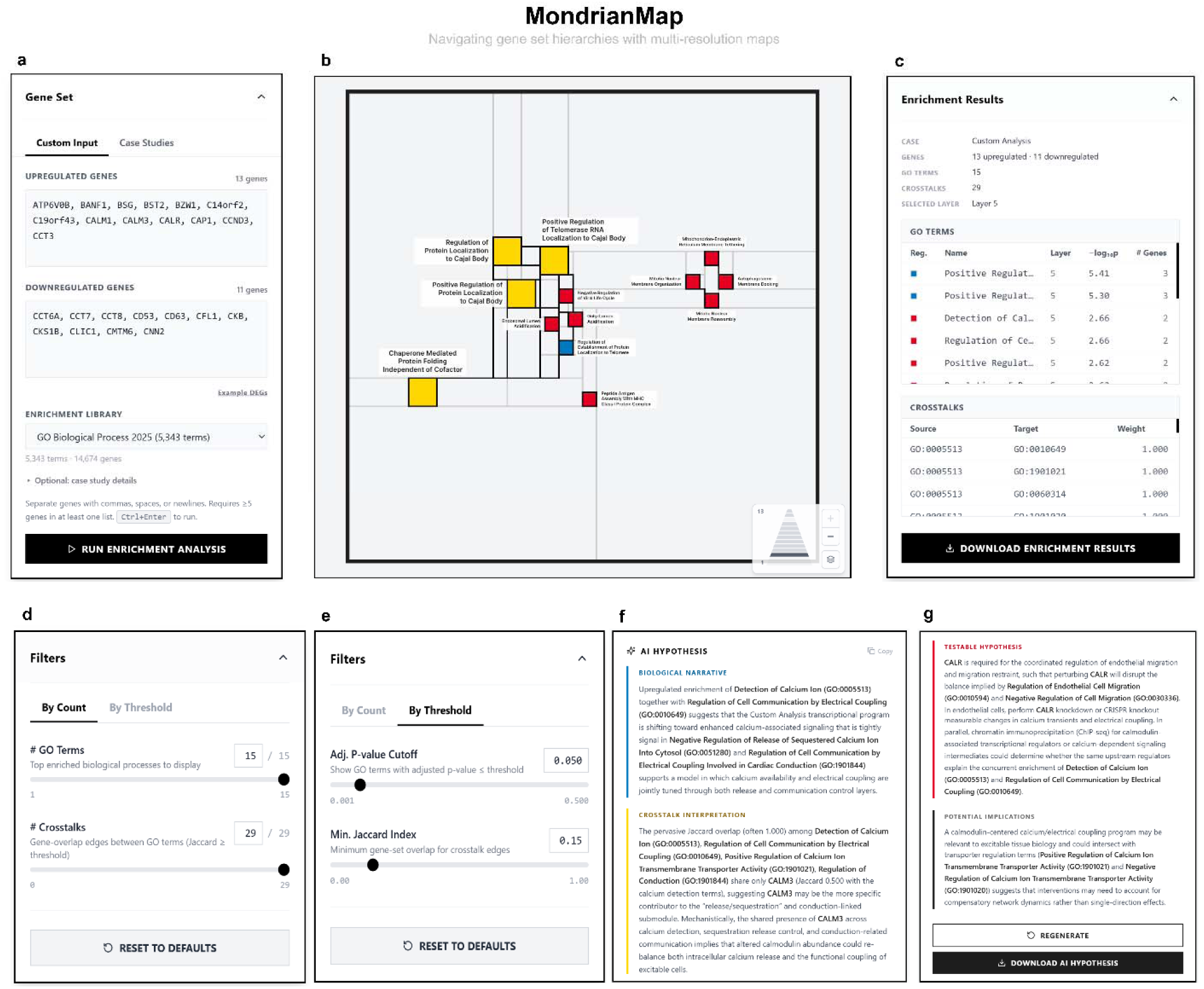
MondrianMap web application interface. (a) Input panel: users select gene sets from curated CFDE collections (LINCS L1000, GTEx, MoTrPAC) or upload custom gene lists for enrichment analysis. (b) Main interactive MondrianMap canvas displaying a hierarchical layout of enriched terms, colored by regulation direction and significance magnitude. GO Terms are hoverable for detailed term information with tooltips. (c) Enrichment results panel showing ranked GO terms with statistics (adjusted p-value, gene count, GOALS layer association). (d–e) Interactive filters control term visibility: by gene count threshold (minimum genes per term) and by statistical significance (adjusted p-value). Likewise, for the crosstalk by count and Jaccard index thresholding. (f–g) The AI hypothesis detail view displays generated hypotheses with supporting genes highlighted in context, enabling rapid validation and interpretation of enrichment patterns.

The AI hypothesis generation module complements the visualization by providing structured narrative summaries grounded in the GOALS layer context. Upon term selection, the module synthesizes a four-part narrative: a. Biological Narrative, b. Crosstalk Interpretation, c. Testable Hypothesis, and d. Potential Implications. Crucially, the LLM is conditioned not only on term-associated genes however, on the GOALS layer context, crosstalks between the terms, and relative position within the biological hierarchy, ensuring that hypotheses directly reflect the enrichment patterns visible in the map itself. This approach extends AI-assisted gene set interpretation to layer-wise, structure-aware summaries. Users should verify all AI-generated claims against the underlying enrichment data and published literature.

### Case Study 1: Hierarchical Visualization Discriminates Cancer Driver Mechanisms

Cancer driver genes converge on overlapping hallmark processes, proliferation, apoptosis evasion, angiogenesis, and immune escape, yet they achieve these outcomes through distinct molecular mechanisms. A conventional enrichment analysis of TP53 knockout versus KRAS knockout in the same cell line will return partially overlapping term lists ranked by p-value, and the investigator is left to infer mechanistic differences by manually comparing hundreds of rows across two spreadsheets. MondrianMap’s layered views resolve this problem visually: by placing two perturbations side by side at a single GOALS layer, mechanistic divergence becomes a visible color contrast. We analyzed LINCS L1000 CRISPR knockout signatures for TP53 and KRAS (two replicates each), all measured in A549 lung adenocarcinoma cells at 96 hours post-perturbation [18], and demonstrate three visual arguments for layer-specific enrichment navigation. A complete summary of enrichment statistics for all analyzed LINCS contrasts is provided in Table S2.

### Cross-Perturbation Contrast at Layer 8: The Same Immune Processes, Opposite Directions

A demonstration of MondrianMap’s capacity for mechanistic discrimination comes from comparing two cancer drivers at a single semantic layer. At GOALS Layer 8, which captures broad cellular-level processes including immune cell migration, differentiation, and metabolic homeostasis, the TP53 and KRAS knockout signatures produce opposite color maps in the same cell line, at the same timepoint, under the same experimental conditions (Figure 3a–b). TP53 knockout at Layer 8 (Figure 3a) shows 13 enriched terms, overwhelmingly downregulated (2 upregulated, 11 downregulated). The blue blocks include Granulocyte Chemotaxis (p = 2.6 × 10⁻³, 8 genes), Neutrophil Chemotaxis (p = 2.9 × 10⁻², 5 genes), Neutrophil Migration (p = 3.8 × 10⁻², 5 genes), Chemokine-Mediated Signaling Pathway (p = 3.5 × 10⁻², 5 genes), Regulation of Blood Coagulation, Cholesterol Homeostasis, and Steroid Metabolic Process. The two red accents are Keratinocyte Differentiation (p = 2.9 × 10⁻L, 8 genes) and Negative Regulation of Striated Muscle Cell Differentiation. The AI hypothesis module identifies two functional hubs: an epithelial stress program driven by ST14, KRT16, S100A7, and SPRR1A/1B (the red), and a suppressed chemokine–coagulation–lipid secretory circuit centered on CXCL10, CXCL9, CXCL5, CCL2, PF4, FGA/FGB/FGG, and APOA2 (the blue). Thus, TP53 loss in A549 cells dampens the tumor’s ability to recruit neutrophils and sustain hemostatic and lipid-transport programs, while activating epithelial differentiation, a phenotypic shift from a secretory, immune-attracting state toward a more differentiated, immunologically quiet state.

**Figure 3.**
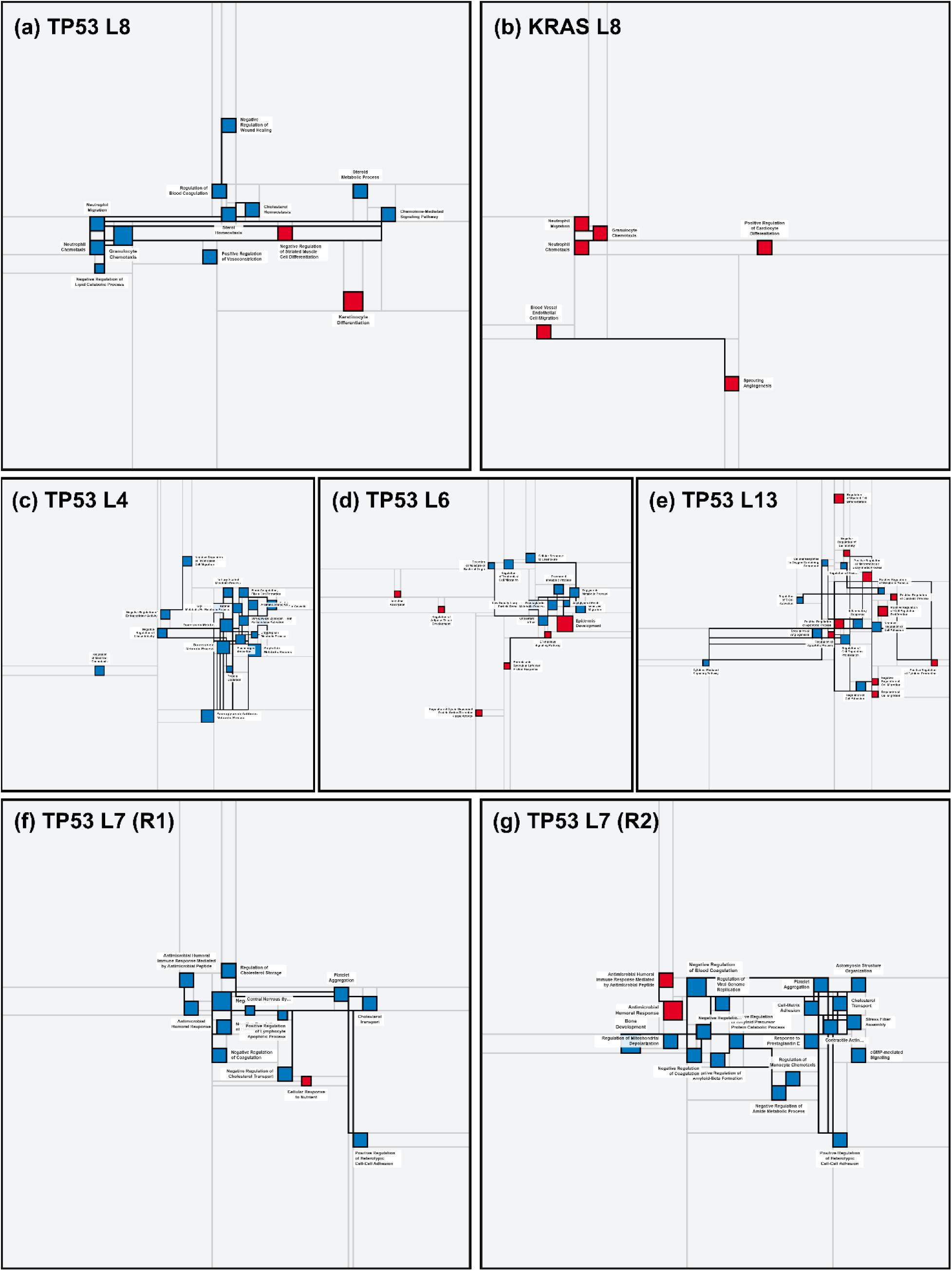
Hierarchical MondrianMaps discriminate cancer driver mechanisms, reveal multi-scale biology, and confirm replicate reproducibility in LINCS CRISPR perturbations. MondrianMap’s layer-specific views expose three dimensions of perturbation biology invisible to flat enrichment outputs. **(a–b)** Cross-perturbation contrast at GOALS Layer 8 (broad cellular processes) between TP53 and KRAS knockouts in A549 lung adenocarcinoma cells: TP53 loss (a) produces a predominantly blue map with suppressed neutrophil chemotaxis, granulocyte chemotaxis, chemokine signaling, and blood coagulation, with small red accents for keratinocyte differentiation; KRAS disruption (b) produces an entirely red map where the *identical* immune cell migration processes, Neutrophil Migration, Granulocyte Chemotaxis, Neutrophil Chemotaxis, are upregulated alongside sprouting angiogenesis and endothelial cell migration, revealing that these two cancer drivers exert *opposite* of these two cancer drivers on the same immune recruitment programs. **(c–e)** Multi-resolution navigation within TP53 knockout across three GOALS layers demonstrates how biological interpretation transforms with semantic resolution: Layer 4 (c) shows a uniformly blue map of 19 downregulated terms dominated by drug metabolism (AKR1C1/C2/C3-driven anthracycline detoxification) and coagulation (fibrinogen); Layer 6 (d) shows an emergent mixed landscape where large red blocks for epithelial differentiation (Epidermis Development, KRT5/14/15) appear alongside persistent blue metabolic suppression (triglycerides, cholesterol, prostaglandins); Layer 13 (e) shows a balanced red-blue mosaic where upregulated apoptosis priming (XBP1, ATF4, ATF6, CDKN2A) and proliferation coexist with downregulated inflammatory response (CXCL10, STAT3) and angiogenesis (VEGFA), illustrating how biological narrative shifts with semantic resolution. **(f–g)** Biological replication validation at GOALS Layer 7 (intermediate immune/defense processes): two independent TP53 knockout replicates produce visually concordant maps, both predominantly blue, both dominated by blood coagulation regulation, antimicrobial humoral response, cholesterol transport, and platelet aggregation, confirming that layer-specific MondrianMaps yield reproducible visual signatures from independent biological replicates.

KRAS knockout at Layer 8 (Figure 3b) is the mirror image: all 6 terms are upregulated, and the map is entirely red. The red blocks are Neutrophil Migration (p = 2.2 × 10⁻², 6 genes), Granulocyte Chemotaxis (p = 4.9 × 10⁻², 6 genes), Neutrophil Chemotaxis (p = 4.9 × 10⁻², 5 genes), the *exact same immune cell recruitment processes* that are blue in the TP53 map, plus Blood Vessel Endothelial Cell Migration (p = 1.3 × 10⁻², 5 genes), Sprouting Angiogenesis (p = 4.1 × 10⁻², 6 genes), and Positive Regulation of Cardiocyte Differentiation. The AI hypothesis module identifies VEGFA, VEGFC, GREM1, and EFNB2 as angiogenic hubs and CXCL8, CXCL6, CXCL5, and S100A8 as chemotactic hubs, interpreting the pattern as a coordinated pro-angiogenic and pro-inflammatory secretion program: KRAS disruption activates compensatory vascular and immune recruitment that the intact KRAS signaling normally suppresses.

The visual comparison is accessible and highlights the directional difference: Neutrophil Chemotaxis, Granulocyte Chemotaxis, and Neutrophil Migration appear as blue blocks in the TP53 map and red blocks in the KRAS map. The same biological processes, at the same semantic layer, in the same cell line, however in opposite directions. This is the mechanistic divergence between two cancer drivers made visible as a color inversion. TP53 loss silences the immune-attracting secretome; KRAS loss activates it. A flat enrichment table would list both perturbations as “significantly enriched for neutrophil chemotaxis” without immediately conveying the directional opposition. The MondrianMaps at Layer 8 make this mechanistic contrast readily apparent.

### Multi-Resolution Navigation: TP53 Knockout Across Layers 4, 6, and 13

The cross-perturbation comparison at Layer 8 establishes that MondrianMap can discriminate between two cancer drivers visually. However, the deeper question is: what does a single perturbation look like when viewed at different semantic resolutions? TP53 knockout provides an ideal test because its 135 enriched terms span all layers from L2 to L13, and the biological narrative transforms dramatically as one traverses the GOALS hierarchy.

At Layer 4 (Figure 3c), the most specific level with substantial enrichment, the MondrianMap shows 19 terms, all downregulated, uniformly blue. The blocks are specific metabolic processes: Aminoglycoside Antibiotic Metabolic Process, Daunorubicin Metabolic Process, Doxorubicin Metabolic Process, Polyketide Metabolic Process (the aldo-keto reductase detoxification pathway), Blood Coagulation Fibrin Clot Formation, Plasminogen Activation, Zymogen Activation, and VLDL Particle Assembly. The AI hypothesis module identifies AKR1C1, AKR1C2, AKR1C3, and AKR1B10 as the metabolic hubs and FGA, FGB, FGG as the coagulation hubs, interpreting this layer as the suppression of a xenobiotic-detoxification and pro-coagulant secretory program. At this resolution, TP53 loss presents as a metabolic perturbation centered on drug metabolism and fibrinogen production.

Ascending to Layer 6 (Figure 3d), the map transforms. Now 16 terms appear, and for the first time, red blocks emerge alongside the blue. The dominant red block is Epidermis Development (p = 4.1 × 10⁻L, 13 genes), driven by keratins KRT5, KRT14, KRT15, SPINK5, and SPRR family proteins, an epithelial differentiation program activated by TP53 loss. Surrounding it are blue blocks: Triglyceride Metabolic Process, Cholesterol Efflux, Prostanoid Metabolic Process, Prostaglandin Metabolic Process, and Low-Density Lipoprotein Particle Remodeling. Additional red accents include Regulation of Adipose Tissue Development, Intestinal Absorption, and ER-Nucleus Signaling Pathway (unfolded protein response). The AI hypothesis module interprets this mixed landscape as TP53 enforcing epithelial differentiation while suppressing lipid handling and eicosanoid metabolism, a less inflammatory, more differentiated state. Layer 6 tells a different story from Layer 4: epithelial remodeling joins the persistent metabolic suppression.

At Layer 13 (Figure 3e), the broadest populated level, the map becomes a balanced mosaic of 20 terms: 10 upregulated (red) and 10 downregulated (blue). The red blocks include Positive Regulation of Apoptotic Process (adjusted p = 5.9 × 10⁻², 15 genes; nominally enriched, above the FDR ≤ 0.05 threshold), Positive Regulation of Cell Population Proliferation (adjusted p = 5.1 × 10⁻², 20 genes; also nominally enriched), Regulation of Myeloid Cell Differentiation, Positive Regulation of Macromolecule Biosynthetic Process, and Positive Regulation of Cytokine Production. The blue blocks include Inflammatory Response (p = 2.5 × 10⁻², 13 genes), Regulation of Angiogenesis (p = 3.3 × 10⁻², 11 genes), Cytokine-Mediated Signaling Pathway, Regulation of Cell Adhesion, and Negative Regulation of Cell Migration. The AI hypothesis module identifies XBP1, ATF4, ATF6, DDIT3, and CDKN2A as the upregulated hubs (UPR and apoptosis priming) and CXCL10, CXCL9, CCL2, STAT3, and VEGFA as the downregulated hubs (inflammatory-angiogenic suppression), interpreting the balanced map as TP53 coupling intrinsic death priming with extrinsic microenvironmental suppression.

The three-layer traversal reveals a principle invisible at any single resolution. Layer 4 says: “TP53 loss suppresses drug metabolism and coagulation.” Layer 6 says: “TP53 loss activates epithelial differentiation while suppressing lipid and prostaglandin pathways.” Layer 13 says: “TP53 loss simultaneously primes the cell for apoptosis and suppresses inflammatory-angiogenic signaling.” These are not contradictions; however, the same perturbation is viewed through progressively broader semantic lenses. MondrianMap’s layer navigation makes this hierarchical logic visible, transforming what would otherwise be a confusing list of 135 terms into a coherent multi-scale biological narrative.

### Biological Replication Consistency: Two TP53 Knockouts at Layer 7

A visualization tool is only useful if it produces consistent results from consistent inputs. Biological replicates, independent CRISPR knockouts of the same gene, should produce visually similar MondrianMaps at the same layer. We tested this with two independent TP53 knockout replicates (K12 and M16) in A549 cells, both measured at 96 hours. At Layer 7 (Figure 3f–g), the two replicate maps share striking visual similarity. TP53 replicate 1 (Figure 3f) shows 13 terms (1 upregulated, 12 downregulated), dominated by blue blocks: Negative Regulation of Blood Coagulation (p = 1.2 × 10⁻³, 7 genes), Antimicrobial Humoral Immune Response (p = 4.9 × 10⁻³, 10 genes), Antimicrobial Humoral Response (p = 6.1 × 10⁻³, 11 genes), Cholesterol Transport, Platelet Aggregation, and Central Nervous System Development. The single red accent is Cellular Response to Nutrient. Hub genes include fibrinogens FGA/FGB/FGG, chemokines CXCL10/CXCL9/CCL2, defensin DEFB1, and apolipoproteins APOA2/APOC1/APOC3.

TP53 replicate 2 (Figure 3g) shows 21 terms (2 upregulated, 19 downregulated), more terms, however, the same visual signature, with overwhelmingly blue with small red accents. Here, we see the same core processes from replicate 1: Negative Regulation of Blood Coagulation (p = 6.6 × 10⁻L, 7 genes), Antimicrobial Humoral Response & Antimicrobial Humoral Immune Response (this time upregulated), Negative Regulation of Coagulation, Platelet Aggregation, and Cholesterol Transport, and Positive Regulation of Heterotypic Cell-Cell Adhesion and the structural similarity confirm that independent TP53 knockouts converge on the same layer-specific functional signature. The additional terms in replicate 2 (actomyosin organization, stress fiber assembly, amyloid regulation) extend, however, do not contradict the replicate 1 pattern, reflecting biological variation at the margins of a reproducible core.

This replicate comparison provides qualitative evidence of consistency, with MondrianMaps from independent knockouts displaying highly concordant visual patterns. Quantitative assessment of replicate concordance (Table S1) further validates this reproducibility. Notably, directional agreement is 100% across 9 of the 12 layers, demonstrating that the dominant visual color per layer is perfectly reproducible. The reduced agreement observed in L7 (75%) and L13 (40%) occurs at the broadest hierarchical levels, where term-level biological stochasticity is expected. Furthermore, L8 exhibits the strongest Spearman correlation (ρ = 0.949, p = 0.051), indicating that quantitative significance magnitudes track closely between replicates at the layer most central to the CS1 cross-perturbation argument. Collectively, these findings confirm that independent knockouts produce robustly concordant maps across comparable layers.

### Case Study 2: Layer-wise MondrianMaps Expose the Inflammaging Duality

Aging is governed by conserved molecular hallmarks, chronic low-grade inflammation, mitochondrial dysfunction, epigenetic drift, and impaired proteostasis [20, 21], yet its manifestation is profoundly tissue-specific. A flat enrichment list for each tissue will report statistically significant GO terms; however, it cannot show the reader whether two tissues age through the same biological process in the same direction, or through the same process in opposite directions, or through entirely different processes. MondrianMaps can answer all three questions in a single visual comparison. We analyzed RNA-seq data from three GTEx tissues, blood, brain, and liver, comparing young adults (20–29 years) to older adults (60–69 years), and additionally compared brain aging at 60–69 with 70–79, where the young group remains the same age at 20–29 years, to assess age-dose progression. Global enrichment statistics and directionality counts for all analyzed tissue and age-group contrasts are detailed in Table S3. The resulting MondrianMaps, viewed at defined GOALS layers, expose the Inflammaging duality as an immediate visual phenomenon and reveal how biological interpretation transforms with semantic resolution and advancing age.

### Cross-Tissue Contrast at Layer 7: The Same Process, Opposite Colors

A clear demonstration of MondrianMap’s cross-tissue discriminative capacity comes from placing three tissues side by side at a single semantic layer. At GOALS Layer 7, which captures intermediate-specificity immune and defense processes, the three aging signatures present radically different color profiles (Figure 4a–c).

**Figure 4.**
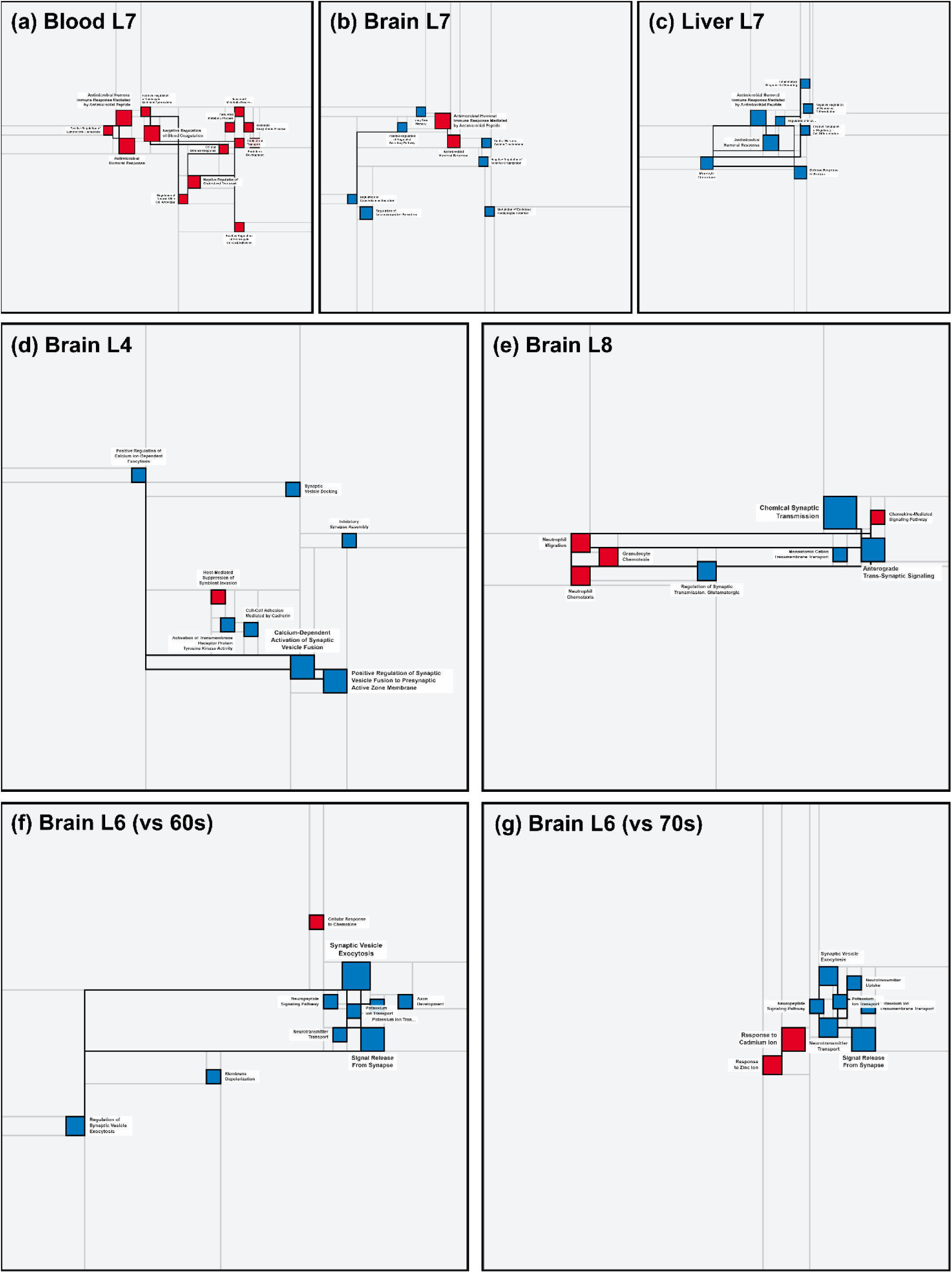
Layer-wise MondrianMaps expose tissue-specific Inflammaging signatures, multi-resolution brain mechanisms, and age-dose progression in GTEx. MondrianMap’s layer-specific views reveal three dimensions of aging biology invisible to flat enrichment outputs. **(a–c)** Cross-tissue comparison at GOALS Layer 7 (intermediate immune/defense processes) across blood, brain, and liver aging (20–29 vs. 60–69 years): blood (a) shows a uniformly red map with 14 upregulated terms dominated by antimicrobial humoral immunity, NK cell activation, and prothrombotic signaling; brain (b) shows a mixed red-blue map where neuroinflammatory chemokine activation (red) coexists with suppression of neurotransmitter secretion, memory, and synapse organization (blue), the tissue-specific dual burden of neurodegeneration and inflammation captured in a single panel; liver (c) shows a uniformly blue map with 8 downregulated terms including the *identical* processes, Antimicrobial Humoral Response, however, in the opposite direction. **(d–e)** Multi-resolution navigation within brain aging (20–29 vs. 60–69) demonstrates how biological interpretation transforms across GOALS layers: Layer 4 (d) shows predominantly blue synaptic machinery loss, calcium-dependent vesicle fusion (SYT1/2/4/7, STX1A), synapse assembly, vesicle docking, with a single red innate immune accent (IFITM1/2/3); Layer 8 (e), the map becomes an evenly balanced red-blue landscape where broad synaptic transmission loss (Chemical Synaptic Transmission, 27 genes) is matched by neutrophil chemotaxis and chemokine signaling activation (CXCL10, CXCL9, S100A8), revealing a neuron-to-neuroimmune state transition invisible at the lower layer. **(f–g)** Age-dose progression in brain at GOALS Layer 6 (intermediate synaptic processes) from the sixties to the seventies: at 60–69 (f), the map is overwhelmingly blue (synaptic vesicle exocytosis, neurotransmitter transport, potassium channel activity) with a single faint red chemokine signal; at 70–79 (g), the blue synaptic decline persists however, two prominent new red blocks emerge, Response to Cadmium Ion and Response to Zinc Ion, driven by the metallothionein gene cluster (MT1M, MT2A, MT1G/H/E/F), marking the decade-specific onset of oxidative metal-stress defense that supplements the earlier inflammatory signature.

Blood aging at Layer 7 (Figure 4a) is uniformly red: all 14 enriched terms are upregulated. The largest blocks are Antimicrobial Humoral Immune Response Mediated by Antimicrobial Peptide (p = 1.5 × 10⁻□, 14 genes), Antimicrobial Humoral Response (p = 1.1 × 10⁻□, 14 genes), and Negative Regulation of Blood Coagulation (p = 2.6 × 10⁻□, 8 genes). The AI hypothesis module identifies KLRC2/KLRC3 (NK cell receptors), DEFA1 (defensin), CXCL8/CXCL9/CCL3/CCL4 (chemokines), and FGA/FGB/FGG (fibrinogen subunits) as hub genes, interpreting the pattern as coordinated antimicrobial-inflammatory activation coupled to prothrombotic hemostatic remodeling, a molecular portrait of Inflammaging in the circulation.

Liver aging at Layer 7 (Figure 4c) is the mirror image: all 8 terms are downregulated, and the map is entirely blue. Remarkably, the two largest blue blocks are the *exact same biological processes* that dominate the blood map, Antimicrobial Humoral Response (p = 2.2 × 10⁻□, 16 genes) and Antimicrobial Humoral Immune Response Mediated by Antimicrobial Peptide (p = 2.2 × 10⁻□, 13 genes), however, in the opposite direction. Alongside them, Defense Response to Fungus, Monocyte Chemotaxis, and Inflammatory Response to Wounding are all suppressed. The AI hypothesis module identifies DEFA4, S100A9, S100A12, IL6, and LTF as hub genes, interpreting this as loss of the liver’s rapid innate inflammatory responsiveness, reduced barrier defense, diminished myeloid recruitment, and attenuated wound repair capacity. This is the Inflammaging paradox made visible: the same defense programs that are *activated* in aging blood are *suppressed* in aging liver. A flat enrichment table would list “Antimicrobial Humoral Response” as significant in both tissues; however, it would not immediately convey the directional inversion. The MondrianMaps at Layer 7 display this relationship as a red-versus-blue contrast for the same GO term.

Brain aging at Layer 7 (Figure 4b) reveals a third pattern: neither purely red nor purely blue, however a mixed landscape of 9 terms (2 upregulated, 7 downregulated). The two red blocks, Antimicrobial Humoral Immune Response and Antimicrobial Humoral Response, echo the blood signature, driven by chemokines CXCL10, CXCL9, CCL4, and CCL19. However, surrounding them are blue blocks: Regulation of Catecholamine Secretion, Positive Regulation of Regulated Secretory Pathway, Long-Term Memory, Regulation of Neurotransmitter Secretion, Central Nervous System Development, Negative Regulation of Synapse Organization, and Modulation of Excitatory Postsynaptic Potential. The AI hypothesis module interprets this coexistence as the brain’s dual aging burden: neuroinflammatory activation (shared with blood) superimposed on progressive synaptic and secretory decline (unique to neural tissue). This bidirectional pattern at Layer 7 is brain-specific.

Together, the three Layer 7 maps provide a visual illustration consistent with the Inflammaging framework: aging activates innate defense in the circulation, suppresses it in the liver, and creates a hybrid state in the brain where inflammation coexists with neurodegeneration. This three-way comparison, difficult to convey through flat enrichment output, is accessible at a glance across three MondrianMaps at the same semantic resolution.

### Multi-Resolution Navigation: Brain Aging at Layer 4 versus Layer 8

The cross-tissue comparison at Layer 7 reveals what the brain shares with and differs from other tissues. However, to understand brain aging *mechanistically*, one must navigate across layers within the brain itself. GOALS Layer 4 captures specific molecular mechanisms, individual signaling events, protein–protein interactions, and precise synaptic processes, whereas Layer 8 captures broader cellular-level processes, synaptic transmission as a category, and immune cell migration as a category. The same tissue at the same age, viewed at these two resolutions, answers fundamentally different questions.

At Layer 4 (Figure 4d), the brain aging map shows 8 terms (1 upregulated, 7 downregulated). The blue blocks are specific synaptic mechanisms: Calcium-Dependent Activation of Synaptic Vesicle Fusion (p = 6.0 × 10⁻□, 5 genes), Positive Regulation of Synaptic Vesicle Fusion to Presynaptic Active Zone Membrane (p = 6.0 × 10⁻□, 5 genes), Inhibitory Synapse Assembly, Synaptic Vesicle Docking, and Positive Regulation of Calcium Ion-Dependent Exocytosis. The AI hypothesis module identifies the synaptotagmin family (SYT1, SYT2, SYT4, SYT7, SYT13) and syntaxin STX1A as hub genes, the calcium-sensing fusion machinery that enables neurotransmitter release. Alongside the synaptic suppression, a single red block appears: Host-Mediated Suppression of Symbiont Invasion, driven by interferon-induced transmembrane proteins IFITM1/2/3, a tiny innate immune signal amid the synaptic decline. This layer tells a story of molecular precision: aging disables the specific protein machinery that executes vesicle fusion at the presynaptic terminal.

Ascending to Layer 8 (Figure 4e), the map transforms. Now the 8 terms divide evenly: 4 upregulated (red) and 4 downregulated (blue), and the visual balance shifts dramatically. The blue blocks are broad synaptic categories, Chemical Synaptic Transmission (p = 4.4 × 10⁻¹¹, 27 genes) and Anterograde Trans-Synaptic Signaling (p = 1.2 × 10⁻□, 18 genes), the aggregate consequence of the Layer 4 fusion machinery loss. However, the red blocks are now equally prominent: Granulocyte Chemotaxis (p = 3.7 × 10⁻□, 9 genes), Neutrophil Chemotaxis (p = 1.6 × 10⁻³, 7 genes), Neutrophil Migration, and Chemokine-Mediated Signaling Pathway. The AI hypothesis module identifies SNAP25, SYT1, GRIN2B, GAD1, and SLC17A7 (glutamate transporter) as the synaptic hubs and CXCL10, CXCL9, CXCL3, CCL4, PF4, S100A8, and CXCR2 as the inflammatory hubs, interpreting this balanced red-blue map as a *neuron-to-neuroimmune state transition*: simultaneous loss of glutamatergic and GABAergic signaling with robust activation of chemokine-mediated neutrophil recruitment into the aging brain.

The two-layer comparison reveals a key insight invisible at either layer alone. Layer 4 shows *what* is breaking: specific calcium-sensing fusion proteins at the presynaptic terminal. Layer 8 shows *what is happening as a consequence*: the aggregate collapse of synaptic transmission coexists with, and may provoke, inflammatory cell infiltration. The tiny red accent at Layer 4 expands into an inflammatory signature at Layer 8, suggesting that the neuroimmune activation is a system-level response to the molecular synaptic failure visible at the lower layer. MondrianMap’s layer navigation makes this mechanistic hierarchy legible in a way that no single-resolution view, whether flat list or aggregated network, could achieve.

### Age-Dose Progression: Brain at Layer 6 from the Sixties to the Seventies

The final dimension of MondrianMap’s discriminative power is temporal, not across tissues or across layers, however across decades within the same tissue at the same semantic resolution. At GOALS Layer 6, which captures intermediate synaptic and signaling processes, brain aging at 60–69 and 70–79 years can be directly compared.

At 60–69 (Figure 4f), the brain L6 map shows 10 terms (1 upregulated, 9 downregulated). The dominant blue blocks are Synaptic Vesicle Exocytosis (p = 9.3 × 10⁻ □, 10 genes), Signal Release From Synapse (p = 1.1 × 10⁻ □, 8 genes), Regulation of Synaptic Vesicle Exocytosis, Neurotransmitter Transport, Potassium Ion Transport, and Membrane Depolarization, a comprehensive suppression of presynaptic function and neuronal excitability. The single red accent is Cellular Response to Chemokine (p = 2.1 × 10⁻², 6 genes), a modest inflammatory signal barely visible against the blue landscape.

At 70–79 (Figure 4g), the same layer shows 9 terms (2 upregulated, 7 downregulated). The blue blocks persist: Signal Release From Synapse, Synaptic Vesicle Exocytosis, Neurotransmitter Transport, Potassium Ion Transport, and Transmembrane Transport remain downregulated. However, the red landscape has changed qualitatively. Two prominent new red blocks have replaced the small chemokine response from the 60s map: Response to Cadmium Ion (p = 7.0 × 10⁻ □, 6 genes) and Response to Zinc Ion (p = 1.2 × 10⁻³, 6 genes). The AI hypothesis module identifies the metallothionein gene cluster, MT1M, MT2A, MT1G, MT1H, MT1E, MT1F, as the hub genes driving these terms, interpreting the shift as the emergence of a metal-buffering and oxidative stress response that was not present at 60–69. Meanwhile, a new blue term appears: Neurotransmitter Uptake (p = 2.6 × 10⁻², 3 genes), driven by SLC32A1 and SLC18A2, indicating that the decade between 60 and 70 extends the synaptic decline from vesicle release (already impaired at 60) to neurotransmitter reuptake machinery.

The visual comparison between the two maps captures the progression of brain aging in a single glance: the blue synaptic loss persists and deepens, while the red component transforms from a faint chemokine signal to a prominent metallothionein stress response. This is not simply “more aging,” however, a qualitative shift in the biology visible at this layer, the emergence of oxidative metal stress as a new dimension of brain aging that supplements (rather than replaces) the inflammatory chemokine signature of the earlier decade. No ranked gene list, p-value histogram, or single-timepoint analysis could convey this decade-over-decade evolution. The layer-specific MondrianMaps place the 60s and 70s side by side and let the color shift speak for itself.

### Case Study 3: Hierarchical Enrichment Reveals Nonlinear Temporal Adaptation Across Tissues

The MoTrPAC Endurance Trained Rats study profiles molecular adaptation to progressive treadmill training at 1, 2, 4, and 8 weeks across multiple tissues [11]. Exercise biology is generally modeled as a gradual, dose-dependent adaptation. MondrianMaps tell a more interesting story: threshold-dependent activation, directional reversals, and tissue-specific biology that only become visible when enrichment is viewed at defined semantic resolutions rather than as aggregated lists. We analyzed brown adipose tissue (BAT) and cardiac ventricle (Heart), two tissues that exhibit contrasting adaptation kinetics, to answer these questions using hierarchical MondrianMaps. A comprehensive summary of enrichment results across all analyzed MoTrPAC tissues and time points is available in Table S4.

### Temporal Threshold: BAT at Layer 3 Reveals a Biological Switch

A clear example of MondrianMap’s layer-specific utility comes from comparing the same tissue at the same semantic layer across time. At GOALS Layer 3, which captures specific molecular mechanisms such as signaling cascades, enzymatic processes, and metabolic reactions, BAT at 2 weeks of training contains just two downregulated terms: L-tryptophan Catabolic Process and Kynurenine Metabolic Process, both driven by KYNU and IDO2 (Figure 5a). The MondrianMap at this layer is nearly empty: two small blue rectangles in a vast white field, indicating that BAT’s early response to exercise is limited to a targeted suppression of immunomodulatory tryptophan catabolism, an early signal of immune quiescence.

**Figure 5.**
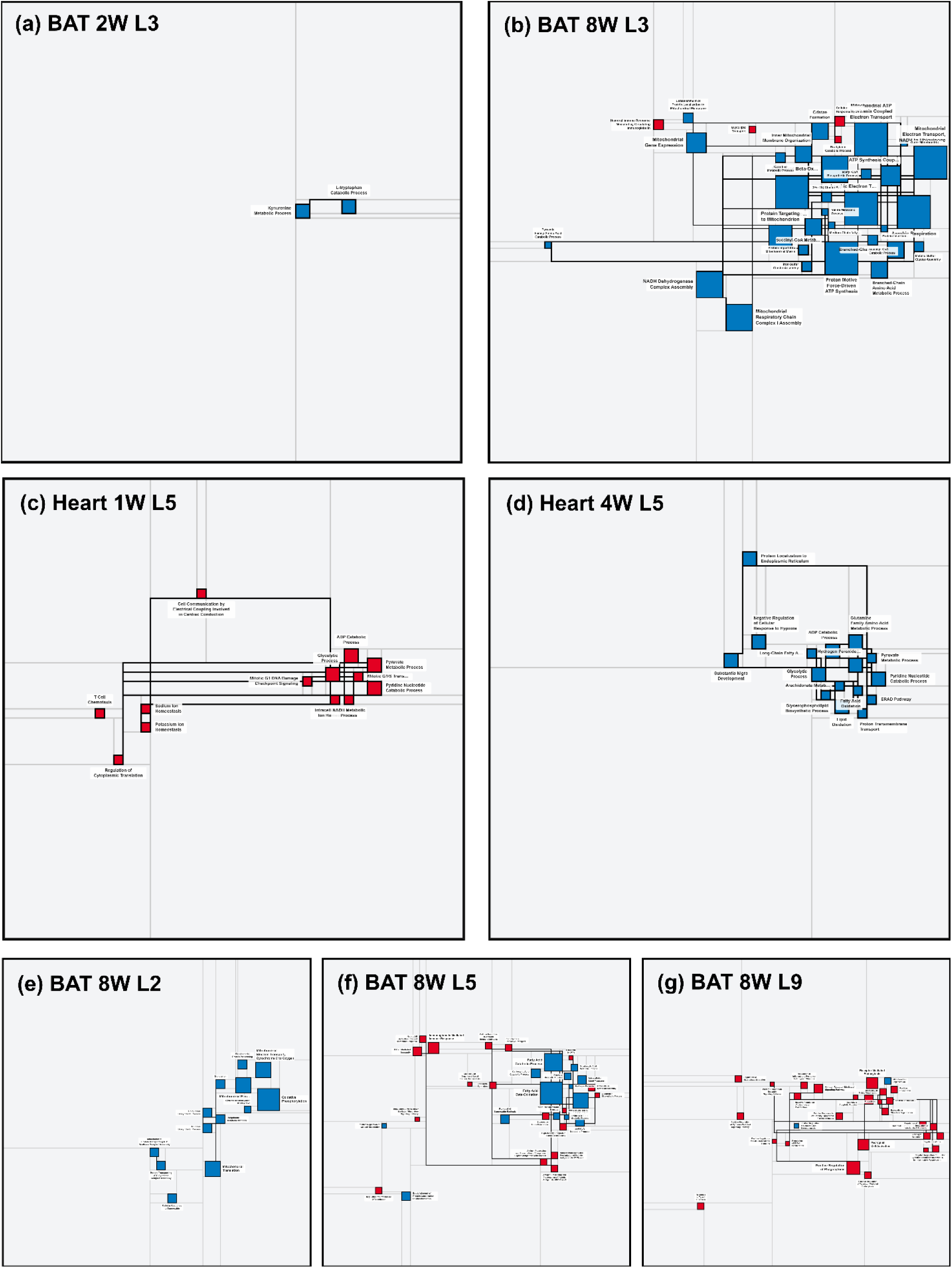
Hierarchical MondrianMaps reveal temporal dynamics and multi-resolution structure in exercise-trained tissues. MondrianMap’s layer-specific views show three dimensions of biological complexity invisible to flat enrichment lists. **(a–b)** Temporal threshold in brown adipose tissue (BAT) at GOALS Layer 3 (specific molecular mechanisms): at 2 weeks of endurance training, only two downregulated kynurenine-pathway terms appear (a); by 8 weeks, the same layer contains 34 terms dominated by mitochondrial respiratory chain assembly, oxidative phosphorylation components, and beta-oxidation, revealing a biological switch from targeted immune modulation to wholesale mitochondrial remodeling (b). **(c–d)** Temporal directional reversal in cardiac ventricle at GOALS Layer 5 (intermediate metabolic processes): at 1 week, all 13 enriched terms are upregulated (red), reflecting acute glycolytic activation and cardiac conduction enhancement driven by PKM, ENO3, and ATP1A3 (c); by 4 weeks, all 16 terms at the same layer are downregulated (blue), indicating suppression of glycolysis, fatty acid oxidation, and ER stress responses through UCP2, ALOX15, and HSPA5, a complete red-to-blue color reversal encoding the transition from acute stress to metabolic efficiency (d). **(e–g)** Multi-resolution navigation within BAT at 8 weeks across three GOALS layers demonstrates how biological interpretation transforms with semantic resolution: Layer 2 (e) shows uniformly downregulated mitochondrial bioenergetics (Oxidative Phosphorylation, p = 9.3 × 10⁻□□); Layer 5 (f) reveals a mixed landscape where metabolic suppression (blue, Fatty Acid Beta-Oxidation) coexists with emerging immune activation (red, B Cell Immunity, Antigen Presentation); Layer 9 (g) is dominated by upregulated innate immune clearance (Phagocytosis, Receptor-Mediated Endocytosis, Microglial Cell Activation), with only residual mitochondrial suppression remaining. Together, these panels demonstrate that layer-wise MondrianMaps resolve temporal switches, directional reversals, and multi-scale biological narratives that collapse into undifferentiated statistics in conventional enrichment outputs.

Six weeks later, the same layer in the same tissue is unrecognizable. BAT at 8 weeks, Layer 3, now contains 34 enriched terms (4 upregulated, 30 downregulated), and the MondrianMap is densely packed with blue rectangles spanning mitochondrial gene expression, inner mitochondrial membrane organization, respiratory chain complex assembly (NADH dehydrogenase, ATP synthase, cytochrome c oxidase), beta-oxidation, and branched-chain amino acid metabolism (Figure 5b). The AI hypothesis module interprets this transition as the onset of comprehensive mitochondrial remodeling: the 2-week tryptophan suppression was an early immunometabolic signal, while the 8-week map reveals wholesale restructuring of the organelle’s translational and bioenergetic machinery. This threshold-like transition is visible at a glance in the layer-specific MondrianMap.

A flat enrichment list would communicate the magnitude, however, not the within-layer mechanistic resolution. A table listing “285 enriched terms at 8 weeks versus 4 at 2 weeks” communicates the magnitude, however, not the mechanistic resolution. The MondrianMap at Layer 3 shows the reader precisely *what* changed and *where in the biological hierarchy* it changed, from two kynurenine-pathway terms to a full mitochondrial rewiring program, within a single visual comparison.

### Temporal Directional Flip: Heart at Layer 5 Shows Adaptation as Color Reversal

The cardiac response to endurance training presents a different kind of temporal complexity: not a threshold explosion, however a directional reversal. At GOALS Layer 5, which encompasses intermediate metabolic and cellular processes, the Heart at 1 week shows 13 enriched terms, all upregulated (Figure 5c). The MondrianMap is populated entirely by red rectangles: ADP Catabolic Process, Glycolytic Process, Pyruvate Metabolic Process, Pyridine Nucleotide Catabolic Process, and Cell Communication by Electrical Coupling Involved in Cardiac Conduction. The AI hypothesis module identifies PKM and ENO3 as the hub genes driving this early glycolytic surge, a coupled metabolic module that enhances ATP generation, redox turnover, and cardiac electrical coupling to meet acute energetic demands. ATP1A3 upregulation reinforces electrogenic ion transport, stabilizing cardiac excitability during the stress of new exercise.

Three weeks later, the same tissue at the same layer has flipped completely. Heart at 4 weeks, Layer 5, shows 16 enriched terms, all downregulated (Figure 5d). The MondrianMap is now entirely blue: Glycolytic Process, ADP Catabolic Process, Fatty Acid Oxidation, Lipid Oxidation, Hydrogen Peroxide Metabolic Process, Protein Localization to Endoplasmic Reticulum, and the ERAD pathway. The AI hypothesis module identifies UCP2, ALOX15, and HSPA5 as the hub genes: mitochondrial uncoupling is suppressed (tighter energetic coupling), lipid-derived inflammatory mediators are attenuated, and ER stress responses are dampened. This coordinated suppression reflects a trained heart that has achieved metabolic efficiency; it no longer needs the stress-buffering and acute energy programs that defined its week-1 response.

This red-to-blue reversal at Layer 5 is the visual signature of cardiac adaptation. A flat enrichment list would show “43 upregulated terms at 1W” and “110 downregulated terms at 4W” as separate, unrelated results. The layer-specific MondrianMaps reveal that the same metabolic processes, glycolysis, ADP catabolism, and pyruvate handling, are first activated and then suppressed, and that this directional flip occurs within a defined semantic stratum. The color reversal between the two maps highlights the transition from acute stress to metabolic efficiency.

### Multi-Resolution Navigation: BAT 8W Across Layers Tells Three Different Stories

The primary demonstration of MondrianMap’s value comes from navigating the same condition across different GOALS layers, revealing how biological interpretation changes with semantic resolution. BAT at 8 weeks provides an ideal test case because its 285 enriched terms span all populated layers (L2–L13), and the biological narrative transforms dramatically as one traverses the hierarchy.

At Layer 2 (Figure 5e), the most mechanistically specific level populated in this dataset, the MondrianMap shows 13 terms, all downregulated. The blue rectangles display Oxidative Phosphorylation (adjusted p = 9.3 × 10□□, 50 genes), Mitochondrial Electron Transport (Complex III → Cytochrome c → Oxygen), Mitochondrial Translation, and ATP Synthase Complex Assembly. This is a map of one story: the systematic suppression of the mitochondrial energy production apparatus. The AI hypothesis module identifies SDHA, SDHB, UQCRC1, UQCRC2, and ATP5F1D as hub genes, interpreting this as coordinated dismantling of the respiratory chain, a remodeling of mitochondrial capacity rather than simple metabolic decline.

Ascending to Layer 5 (Figure 5f), the map transforms. Now 32 terms appear, and for the first time, red and blue rectangles coexist. The dominant blue blocks are Fatty Acid Beta-Oxidation (p = 4.8 × 10⁻²², 23 genes) and Fatty Acid Catabolic Process, continued metabolic suppression, consistent with Layer 2. However, alongside them, red blocks emerge: Immunoglobulin Mediated Immune Response, B Cell Mediated Immunity, Antigen Processing and Presentation via MHC Class I and Class II, and Extracellular Matrix Disassembly. The hub genes shift from mitochondrial subunits to immune effectors (CD74, CD19, CTSS, CTSL) and tissue remodelers (MMP2, MMP3, MMP12, MMP14). The AI hypothesis module interprets this mixed landscape as a metabolic-to-immune pivot: lipid catabolism is suppressed while adaptive immunity and tissue remodeling are activated, a shift invisible at Layer 2, where only the mitochondrial story exists.

At Layer 9 (Figure 5g), the map shifts again, now dominated by upregulated terms. Of 29 terms, 27 are red. The largest blocks are Positive Regulation of Phagocytosis (p = 1.2 × 10□, 15 genes), Receptor-Mediated Endocytosis, Microglial Cell Activation, and Antigen Receptor-Mediated Signaling Pathway. The hub genes are innate immune receptors and effectors: FCER1G, FCGR1A, CLEC7A, MSR1, MRC1, ITGAM, ITGB2, TYROBP, and complement components C3, C1QA, C5AR1. The AI hypothesis module characterizes this as a dominant innate immune clearance program, phagocytic remodeling with complement activation, and macrophage-like cell recruitment. The only blue block is Mitochondrion Organization (12 genes), a residual echo of the mitochondrial suppression story from Layer 2, now marginalized by the overwhelming immune activation.

The three-layer traversal tells a story that no single flat list, ranked table, or aggregated visualization could convey. Layer 2 says: “BAT is dismantling its mitochondrial machinery,” Layer 5 says: “BAT is simultaneously suppressing fat oxidation and activating adaptive immunity,” and Layer 9 says: “BAT is recruiting innate immune cells for tissue remodeling.” These are not three different analyses; they are three views of the same biological event, resolved at different semantic scales. MondrianMap’s layer navigation makes this multi-scale biology legible in a way that collapses into noise in any flat representation.

### Tissue Contrast: Same Layer, Different Tissues, Different Regulations

The layer-specific view also enables direct cross-tissue comparison at matched semantic resolution. At Layer 5, BAT at 8 weeks (Figure 5f) shows a mixed red-blue landscape (metabolic suppression plus immune activation), while Heart at 4 weeks (Figure 5d) shows pure blue (metabolic efficiency). Both tissues are undergoing exercise-induced remodeling; however, the color patterns at the same layer reveal fundamentally different biological strategies: BAT recruits immune surveillance while suppressing metabolism, whereas the heart simply quiets its metabolic stress responses. This tissue-level contrast, visible as a color difference between two MondrianMaps at the same layer, directly demonstrates that exercise adaptation is not a single program applied uniformly across tissues; however, a set of tissue-specific strategies operating within shared semantic categories.

## DISCUSSION

MondrianMap addresses a fundamental gap in how biologists interpret enrichment results: existing tools can rank terms by statistical significance, cluster them by semantic similarity, or connect them by gene overlap, however, none can show the reader what happens to a biological process when you change the tissue, the timepoint, the perturbation, or the level of abstraction, all within a single visual framework. The shift from flat ranked lists to layer-specific, color-encoded rectangular maps provides a visual reasoning framework that complements statistical enrichment analysis. Across three case studies spanning cancer pharmacogenomics, aging biology, and exercise physiology, MondrianMap facilitated biological comparisons that are cumbersome to perform with flat outputs.

### Layer-Specific Navigation Exposes Mechanistic Logic Hidden in Flat Lists

The central proposition of MondrianMap is that viewing enrichment at defined GOALS layers facilitates recognition of patterns that are difficult to discern in aggregated views. Our three case studies provide converging evidence for this claim, each demonstrating a different facet of what layer resolution makes visible.

In the LINCS case study, the critical insight emerged not from examining a full GO term list, but from placing TP53-KO and KRAS-KO side by side at a single layer. At GOALS Layer 8, three immune cell migration processes, Neutrophil Chemotaxis, Granulocyte Chemotaxis, and Neutrophil Migration, appear as blue blocks in the TP53 map and red blocks in the KRAS map (Figure 3a–b). The same biological processes, in the same cell line, at the same timepoint, with opposite regulations. A ranked enrichment table would report both perturbations as “significantly enriched for neutrophil chemotaxis” and leave the investigator to discover the directional inversion by cross-referencing fold-change columns. The MondrianMaps make the opposition self-evident: blue versus red for the same rectangular block. This is not a subtle statistical finding; It is a visual pattern that highlights the contrasting tumor microenvironmental consequences of TP53 loss (immune silencing) versus KRAS disruption (immune and angiogenic activation).

The TP53 multi-resolution traversal deepened this insight. At Layer 4, the map was uniformly blue, drug metabolism and coagulation factor suppression, a story about xenobiotic detoxification (AKR1C family), and fibrinogen loss. At Layer 6, red blocks emerged: Epidermis Development (KRT5/14/15) activated alongside persistent metabolic suppression. At Layer 13, the map showed a red-blue mosaic in which apoptosis-related and inflammatory terms appeared, though two terms (Positive Regulation of Apoptotic Process and Positive Regulation of Cell Population Proliferation) had adjusted p-values marginally above the 0.05 threshold (5.9 × 10□² and 5.1 × 10□², respectively) and should be interpreted with caution. The three layers told three different, however, internally consistent stories: specific metabolic rewiring, intermediate epithelial-metabolic remodeling, and system-level coupling of death readiness with microenvironmental quieting (Figure 3c–e). MondrianMap’s layer system imposed the structure that made the narrative legible.

The GTEx aging case study demonstrated that the same layer-specific logic applies across tissues. At Layer 7, the Inflammaging duality became a color photograph: Blood showed 14 red blocks (antimicrobial immunity, NK cell activation, coagulation), Liver showed 8 blue blocks (the *same* Antimicrobial Humoral Response process, however, suppressed), and Brain showed a mixed red-blue map where neuroinflammatory chemokines coexisted with suppressed catecholamine secretion and synapse organization (Figure 4a–c). The directional inversion between Blood and Liver at Layer 7, identical GO terms in opposite colors, is the Inflammaging paradox rendered visible. A conventional analysis reporting “antimicrobial response significant in both tissues” would miss the opposition entirely. MondrianMap’s directional color-encoding makes this contrast readily apparent.

The Brain multi-resolution comparison added mechanistic depth. At Layer 4, the map was almost entirely blue, with specific synaptic fusion machinery (SYT1/2/4/7, STX1A) breaking down. At Layer 8, the map became an evenly balanced red-blue landscape where broad synaptic transmission loss met neutrophil chemotaxis and chemokine signaling activation (Figure 4d–e). The tiny red immune accent at Layer 4 expanded into a full inflammatory signature at Layer 8, revealing a neuron-to-neuroimmune state transition that unfolds across the semantic hierarchy. The age-dose comparison at Layer 6 further showed that the red component evolved qualitatively between decades: a faint chemokine signal at 60–69 was replaced by prominent metallothionein-driven metal-stress blocks (MT1M, MT2A, MT1G/H/E/F) at 70–79 (Figure 4f–g), not simply “more aging"; however, the emergence of a new biological dimension.

The MoTrPAC case study demonstrated that layer resolution captures temporal dynamics. Comparing BAT at Layer 3 across 2 and 8 weeks of training revealed a biological switch: two small blue kynurenine-pathway blocks at 2 weeks became 34 dense blue blocks of mitochondrial respiratory chain assembly at 8 weeks (Figure 5a–b). The Heart at Layer 5 showed a complete red-to-blue directional reversal: all 13 terms upregulated (glycolytic activation, PKM/ENO3) at 1 week, all 16 terms downregulated (metabolic suppression, UCP2/ALOX15/HSPA5) at 4 weeks (Figure 5c–d). And navigating BAT at 8 weeks across Layers 2, 5, and 9 revealed a story that transformed at every resolution: uniformly blue mitochondrial dismantling (L2), mixed metabolic suppression plus immune activation (L5), predominantly red innate immune clearance with phagocytosis and complement activation (L9) (Figure 5e–g). These temporal and multi-resolution patterns, threshold switches, directional reversals, and layer-dependent narrative shifts are precisely the dynamics that flat enrichment tools were never designed to capture.

### Directional Color-Encoding as a First-Class Visual Variable

A limitation common to all significance-only visualization tools is the loss of effect direction when terms are grouped, clustered, or hierarchically organized. By encoding effect direction independently from significance, red for upregulated genes, blue for downregulated, MondrianMap treats directionality as a first-class visual variable rather than an annotation buried in a supplementary column.

This design choice proved consequential across all three case studies. The Inflammaging duality in GTEx, up-immune in Blood, down-metabolic in Liver, was visible as a red-versus-blue contrast at Layer 7 without requiring any statistical comparison. The TP53-versus-KRAS mechanistic divergence at Layer 8 was visible as a blue-versus-red inversion of the same GO term blocks. The Heart exercise adaptation from week 1 to week 4 was visible as a complete color flip at Layer 5. In each case, the biological insight was encoded in color, not in a p-value table, and the reader could perceive the finding before consciously interpreting the individual terms. This immediacy, the ability to see a directional pattern before reading the labels, is what distinguishes a visualization tool from a statistical report, and it is what MondrianMap’s color scheme was designed to achieve.

### Comparison to Prior Art: Why Layer Structure Matters

How does MondrianMap compare to existing tools designed to handle enrichment complexity? REVIGO [12] pioneered semantic similarity clustering, reducing large term sets to non-redundant representatives plotted in two dimensions. This is effective for small enrichment sets; however, the 2D plane collapses hierarchical depth, and its static output does not support multi-resolution navigation. Enrichment Map [13], implemented in Cytoscape, constructs networks where nodes are gene sets and edges reflect gene overlap, effective at a moderate scale; however, increasingly cluttered for large enrichment sets. Neither REVIGO nor Enrichment Map encodes effect direction as a primary visual channel.

The treemap and sunburst approaches implemented in GO browsers faithfully represent the ontology’s graph structure; however, they cannot simultaneously encode enrichment statistics, effect direction, and semantic clustering without visual overload. clusterProfiler [14], a widely used R package, offers semantic similarity grouping and ridge plots for multi-contrast comparison; however, relies on ranked lists and lacks the explicit, quantitative layer stratification that GOALS provides. Its groupings are data-driven clusters rather than biologically principled strata, and it cannot show the reader what happens to the same enrichment when viewed at a different semantic resolution.

MondrianMap’s advantage lies not in any single feature however, in its combination within a principled hierarchical framework. The GOALS layer system, grounded in the co-membership geometry of the GO graph, with power-law-distrihowever,ed term assignments (R² = 0.98) and two orders of magnitude variance reduction compared to depth-based stratification [15], provides a biological scaffold that semantic similarity alone cannot match. Layers are not ad-hoc clusters; they are quantitatively defined strata of biological abstraction. This is what enables the multi-resolution traversal that proved essential in all three case studies: the ability to ask “what does this perturbation look like at the level of specific enzyme reactions?” and then ask “what does it look like at the level of system-wide regulation?” and receive coherent, non-contradictory answers that build on each other.

### Crosstalk, Hub Genes, and AI-Guided Hypothesis Generation

Beyond visualization, MondrianMap’s Jaccard Index-based crosstalks and integrated AI hypothesis module add interpretive layers that contextualize what the maps display. Terms sharing genes form crosstalk clusters visible as connected blocks within each layer view; the genes driving these connections, identified by the AI module, are candidate integrators for experimental follow-up. In the LINCS case, the AI module traced the TP53 Layer 13 balanced mosaic to specific hub genes: XBP1, ATF4, and ATF6 coupling unfolded protein response to apoptosis priming (red), while CXCL10, STAT3, and VEGFA coordinated the suppressed inflammatory-angiogenic axis (blue). In the MoTrPAC BAT 8W Layer 9, the module identified FCER1G, CLEC7A, TYROBP, and CSF1R as innate immune clearance hubs linking phagocytosis to complement activation. These narratives emerged directly from the layer-specific enrichment structure and were immediately verifiable against the visible map, a key distinction from unconstrained LLM summaries that lack grounding in hierarchical context.

The AI hypothesis module is best understood as an optional structured reading aid for the MondrianMaps, *not as an autonomous discovery engine*. Its four-part output, Biological Narrative, Crosstalk Interpretation, Testable Hypothesis, and Potential Implications, translates the visual patterns of each layer into the language of experimental biology. The grounding in the GOALS layer context ensures that hypotheses reflect the semantic resolution at which they were generated: a Layer 4 hypothesis discusses specific enzymatic mechanisms, whereas a Layer 13 hypothesis discusses system-level regulatory programs. This resolution-awareness distinguishes MondrianMap’s AI module from general-purpose gene list summarizers and anchors its outputs in the hierarchical structure that defines the tool.

### Cross-Dataset Integration and the Exercise-Reverses-Aging Hypothesis

A major advantage of CFDE integration is the ability to compare enrichment patterns across independent, large-scale programs. The GTEx aging and MoTrPAC exercise response cases, viewed through layer-specific MondrianMaps, provide a visual basis for the hypothesis that exercise counteracts aging transcriptomic signatures. The Inflammaging duality observed in GTEx, upregulated immune processes (red) in Blood, downregulated metabolic processes (blue) in Liver at Layer 7, is precisely inverse to the exercise response patterns in MoTrPAC. BAT at 8 weeks shows upregulated immune remodeling (red at Layer 9) alongside downregulated mitochondrial and metabolic programs (blue at Layer 2), while the Heart at 4 weeks shows broad metabolic suppression (all blue at Layer 5) following an initial activation phase (all red at Layer 5, week 1). The directional patterns suggest that exercise and aging engage the same ontological strata; however, it push them in opposing directions, a hypothesis that becomes visually tractable only when enrichment is resolved by layer and encoded by color.

MondrianMap’s CFDE integration provides reusable community value beyond the three case studies presented here. The pre-indexed LINCS, GTEx, and MoTrPAC enrichment repositories are accessible to any user without requiring local computation or data wrangling, enabling researchers to perform cross-program comparisons at matched semantic resolution. This interoperability exemplifies the CFDE vision of making Common Fund datasets more accessible: a user studying exercise biology in MoTrPAC can compare layer-specific patterns against GTEx aging signatures in seconds, a comparison that would otherwise require downloading, re-processing, and aligning two independent datasets.

While direct temporal coordination across these independent studies is not possible, MondrianMap’s layer-stratified views provide the framework for systematic comparison. Future work integrating longitudinal aging cohorts with exercise intervention data could formally test this reversal hypothesis at the systems biology level, tracking layer-by-layer signature evolution to determine which layers respond earliest, which resist intervention, and which semantic strata harbor the molecular targets most amenable to exercise-based or pharmacological reversal of aging phenotypes.

### Limitations and Considerations

While MondrianMap effectively bridges mechanistic detail and system-level themes for cross-condition and cross-tissue comparisons, several limitations and potential pitfalls warrant careful consideration to ensure robust interpretation. The tool is most informative when input gene sets possess sufficient moderate-to-large effect sizes to populate multiple GOALS layers, typically requiring 50 or more enriched terms. Extremely sparse enrichments of fewer than 5 terms per layer, often seen in datasets with low transcriptomic change, produce visually uninformative maps and are better analyzed using conventional tables or dot plots. Conversely, highly dense enrichments exceeding 300 terms per layer require significance or gene-count filtering to maintain visual interpretability, a capability supported within the web interface. Furthermore, the framework is currently restricted to the GO Biological Process ontology to maintain simplicity in study design. Extending the GOALS hierarchical framework to pathway databases such as KEGG [27], Reactome [28], and MSigDB [29] remains a priority for future development.

Users must also exercise caution when interpreting the spatial and color encodings of the generated maps. Adjacency in MondrianMap reflects GoBERT embedding similarity projected via UMAP rather than direct ontological parent–child relationships, meaning spatial proximity should not be misinterpreted as strict hierarchical ancestry. While the UMAP projection is seeded for reproducibility, its stochastic nature means minor layout variations may still occur across different software versions. Additionally, color gradients are inherited directly from upstream differential expression analyses. Because different pipelines, such as Characteristic Direction for LINCS versus log-fold-change for GTEx, can assign conflicting directionality to the exact same biological process, cross-dataset color comparisons can be complicated. Analytically, enrichments captured at the broadest GOALS layers may include terms with adjusted p-values near the significance threshold, and these should be interpreted as suggestive system-level themes rather than definitive mechanisms.

Finally, the integrated AI hypothesis module is an optional, assistive feature rather than a validated core contribution. Its outputs are heavily conditioned on the underlying large language model’s training data, prompt structure, and deployed model version. The module can occasionally produce confident-sounding narratives even when the underlying enrichment data is sparse or ambiguous. Because a systematic evaluation of hallucination rates, citation accuracy, and expert agreement has not yet been conducted, users must treat all AI-generated hypotheses as speculative starting points that require rigorous manual verification against raw enrichment statistics and published literature. A small-scale expert audit of AI outputs across representative layers is a priority for future work to strengthen confidence in this module.

### Future Directions

MondrianMap’s development points toward several natural extensions. We envision multi-contrast comparison views that directly overlay MondrianMaps for two or more conditions at the same layer, enabling the side-by-side visual comparisons demonstrated across our case studies to be performed interactively for arbitrary pairs of contrasts. Extension of the GOALS hierarchical framework to non-GO gene set libraries would broaden applicability to mechanistic pathway databases and disease-specific signatures. A programmatic API would further enable integration into scripted pipelines, supporting bulk enrichment comparison and automated hypothesis generation across large cohorts of experimental contrasts. Together, these directions would position MondrianMap not merely as a visualization endpoint, however, as an active research instrument for layered, cross-dataset biological discovery.

## METHODS

### GO Enrichment Analysis

Gene set enrichment analysis was performed using Fisher’s exact test with Benjamini–Hochberg false discovery rate (FDR) correction, with adjusted p-value threshold p ≤ 0.05. The background gene universe was defined as the union of all unique genes across the reference gene set library. The reference gene set library used was GO_Biological_Process_2025. For LINCS contrasts, the top 250 upregulated genes and top 250 downregulated genes, ranked by Characteristic Direction magnitude [6], were analyzed as input sets. The Characteristic Direction method estimates directionality of gene expression change robust to noise and batch effects; genes with the highest absolute Characteristic Direction scores represent the most significant directional shift in the contrast. Enriched terms derived from upregulated genes were labeled as “upregulated” (visualized in red), those from downregulated genes as “downregulated” (blue), and terms appearing in both sets as “shared” (yellow).

For GTEx and MoTrPAC contrasts, pre-computed differential expression signatures from the respective CFDE databases were used as input. Directional labeling (red for upregulated, blue for downregulated) was assigned based on the sign of the logL fold-change of the leading edge genes in each enriched term.

### GO Layer Assignment

Enriched GO-BP terms were assigned to semantic layers L1–L13 using the GOALS framework [15]. GOALS applies an Initiation-Assignment-Expansion-Repetition (IAER) procedure to the GO Biological Process co-membership network. The algorithm iteratively assigns terms to layers such that the resulting layer-wise distribution of term sizes achieves a power-law fit with exponent β and R² = 0.98, while simultaneously reducing the mean sample variance of gene set sizes by approximately two orders of magnitude compared to conventional GO depth-based stratification. GO layer assignments for GO_Biological_Process_2025 were pre-computed and are provided as a CSV reference file in the MondrianMap codebase.

### Semantic Layout and Embedding-Based Positioning

Overall, GO terms are positioned in two-dimensional space using GoBERT embeddings [30] combined with UMAP dimensionality reduction. GoBERT is a BERT-based language model fine-tuned on the full corpus of GO term descriptions, producing high-dimensional dense vectors (1024 dimensions) that encode semantic relationships between terms as inferred from their textual definitions and position in the GO graph. These high-dimensional vectors are projected to two dimensions using Uniform Manifold Approximation and Projection (UMAP) [31] with parameters: n_neighbors = 15 (dynamically adjusted to min(n_neighbors, n_terms − 1), floored at 2), min_dist = 0.1, metric = cosine, random_state = 42, n_components = 2. These parameters preserve local neighborhood structure such that terms with high embedding similarity are positioned in spatial proximity. However, while UMAP is inherently stochastic at smaller sample sizes, quantitative assessment confirms that the local semantic clustering and hierarchy-tier segregation remain highly stable across random seeds, even when global orientations vary (Figure S2).

Block rectangles are positioned according to the UMAP coordinates of their corresponding GO terms. Block area is proportional to −log□□ (adjusted p-value), encoding effect magnitude as visual weight. Blocks are colored by enrichment direction: red for upregulated, blue for downregulated, and yellow for shared between up and down. The rectangular decomposition uses a treemap-style algorithm [32] to tile the available space without overlap while preserving the approximate position of each term’s centroid.

### Crosstalk Edge Computation

Pairwise Jaccard indices were computed between all enriched GO term gene sets within each contrast using the standard formula: Jaccard(A, B) = |A ∩ B| / |A L B|. Term pairs with Jaccard index ≥ 0.15 are connected by visible crosstalk edges in the interactive MondrianMap visualization, by default. Edge width is proportional to the Jaccard coefficient, such that high-overlap term pairs are visually emphasized as thick edges. The threshold of 0.15 was chosen to balance edge visibility with information density. To evaluate how this parameter shapes the network topology that underpins our spatial layout, we assessed network characteristics across Jaccard thresholds from 0.05 to 0.30 (Figure S1). The 0.15 default occupies an optimal transition zone: it successfully prunes spurious, low-overlap edges while preserving the global connectivity (retaining 97% of terms in connected components) required for UMAP to embed semantically related terms nearby. Users may adjust this parameter in the web interface to suit specific analytical requirements.

### Reproducibility Parameters

GO Library: GO_Biological_Process_2025. Background Universe: union of all genes in the reference GMT file. Enrichment Test: scipy.stats.fisher_exact (one-tailed, alternative=“greater”). Multiple Testing: Benjamini–Hochberg FDR, threshold adjusted p ≤ 0.05. GoBERT Model: MM-YY-WW/GoBERT (HuggingFace), 1024-dimensional embeddings. UMAP: umap-learn ≥ 0.5.3; n_neighbors=15, min_dist=0.1, metric=cosine, random_state=42, n_components=2. Jaccard Threshold: 0.15 (default; user-adjustable in interface). AI Module: OpenAI API, temperature=0.1, max_completion_tokens=1500, model=gpt-5.4. Key Python Dependencies: pandas ≥ 2.0.0, numpy ≥ 1.24.0, scipy ≥ 1.10.0, torch ≥ 2.0.0, transformers ≥ 4.30.0, scikit-learn ≥ 1.3.0, flask ≥ 3.0.0. Frontend: React 18.3.1, D3 7.9.0, Vite 5.3.1. Source Code: https://github.com/aimed-lab/mondrian-web.

### AI Hypothesis Generation Module

The AI hypothesis module accepts as input: (i) the list of significantly enriched GO terms at user-selected layers; (ii) the gene members of each term; (iii) the pairwise Jaccard crosstalk matrix; and (iv) the layer context from GOALS. These data are formatted into a structured prompt and passed to the OpenAI API (model=gpt-5.4; temperature = 0.1; max_completion_tokens = 1,500). The system prompt instructs the model to act as a “computational biology expert and gene set analyst,” reference every GO term and gene symbol exactly as provided, and never fabricate identifiers or p-values. The prompt elicits four specific sections: (1) Biological Narrative, a synthesis of the enrichment pattern, interpreting upregulation and downregulation in biological context; (2) Crosstalk Interpretation, explanation of the hub genes and functional modules revealed by the crosstalk network; (3) Testable Hypothesis, a specific, mechanistic hypothesis that could be evaluated experimentally; and (4) Potential Implications, discussion of the findings’ relevance to disease biology, therapeutic targets, or broader biological principles. Rate limiting (100 requests per hour per session) and explicit grounding constraints (exact gene symbols, exact GO identifiers, no fabricated statistics) serve as the primary guardrails against hallucination. Users are instructed to treat AI-generated hypotheses as speculative and to validate all claims against the peer-reviewed literature and, ideally, through experimental evidence.

### CFDE Database Integration

MondrianMap integrates three publicly accessible CFDE program databases as pre-indexed, searchable repositories. No new experiments or human subjects research were conducted for this study.

1. **LINCS L1000 CRISPR Perturbation.** Genome-wide transcriptomic (L1000) measurements in multiple cell lines following CRISPR knockout or overexpression of individual genes [6, 7]. Users select a gene and cell line to retrieve baseline contrasts and pre-computed enriched gene sets.
2. **GTEx Aging Signatures 2021.** Differential expression signatures comparing old (≥60 years) versus young (20–40 years) individuals across 48 tissues, derived from the GTEx project [8, 9]. Pre-computed enriched gene sets for each tissue are accessible within MondrianMap.
3. **MoTrPAC Endurance Trained Rats 2023.** Transcriptomic time-course data from rats subjected to endurance treadmill training across 19 tissues at 1, 2, 4, and 8 weeks [10, 11]. Original animal care and use were approved by the respective institutions as reported in the primary publications. Pre-computed enriched gene sets are available for each tissue and time point.

### Web Application Architecture

MondrianMap is implemented as a React.js single-page application (SPA) with a Python backend. The frontend is a responsive interface enabling interactive exploration of MondrianMaps: users can zoom in/out to inspect specific GO layers and terms, hover to see term definitions and member genes, and click to view crosstalk edges and launch the AI hypothesis module. The backend performs GO enrichment computation (Fisher’s exact test, FDR correction) and interfaces with the large language model API for hypothesis generation. The application is deployed at https://mondrianmap.smartdrugdiscovery.org/. Source code is available at https://github.com/aimed-lab/mondrian-web under an open-source license.

## RESOURCE AVAILABILITY

### Lead Contact

Further information and requests for resources and reagents should be directed to and will be fulfilled by the Lead Contact, Jake Y. Chen.

### Materials Availability

This study did not generate new, unique reagents or materials. All external data sources and databases used are cited in the manuscript.

### Data and Code Availability

MondrianMap source code is available at https://github.com/aimed-lab/mondrian-web. The web application is deployed and freely accessible at https://mondrianmap.smartdrugdiscovery.org/. Pre-computed enrichment results for all case studies and GO layer assignments are available through the MondrianMap interface and associated GitHub repository.

## ACKNOWLEDGEMENT

This work was supported by a grant from the National Institute of Health (Award No. U54:OD036472) and the startup fund (PI: Jake Y. Chen) of System Pharmacology AI Research Center at the University of Alabama at Birmingham. The authors acknowledge the National Artificial Intelligence Research Resource (NAIRR) Pilot and OpenAI API credits for contributing to this research result. We also thank the NIH Common Fund programs LINCS, GTEx, and MoTrPAC for making their datasets publicly available through the Common Fund Data Ecosystem (CFDE).

## AUTHOR CONTRIBUTIONS

F.A. is the lead author and primary developer of the project, executing the core software implementation and writing the original draft of the manuscript. Z.Y. provided the foundational GOALS hierarchical framework, which served as the primary data structure for the hierarchical representation. E.S. designed and implemented the data processing pipeline. M.D.H. developed the semantic layout algorithms. Z.S. built and managed the system infrastructure and deployment. S.Z. provided project supervision and significantly contributed to the writing, reviewing, and editing of the manuscript. J.Y.C. conceived the study, acquired funding, supervised all aspects of the research, and contributed to the writing and critical revision of the paper. All authors participated in reviewing and editing the final manuscript.

## REFERENCES

1. Chen, E.Y., Tan, C.M., Kou, Y., Duan, Q., Wang, Z., Meirelles, G.V., Clark, N.R., & Ma’ayan, A. (2013). Enrichr: interactive and collaborative HTML5 gene list enrichment analysis tool. BMC Bioinformatics, 14, 128. 10.1186/1471-2105-14-128

2. Kuleshov, M.V., Jones, M.R., Rouillard, A.D., Fernandez, N.F., Duan, Q., Wang, Z., Koplev, S., Jenkins, S.L., Jagodnik, K.M., Lachmann, A., McDermott, M.G., Monteiro, C.D., Gundersen, G.W., & Ma’ayan, A. (2016). Enrichr: a comprehensive gene set enrichment analysis web server 2016 update. Nucleic Acids Research, 44(W1), W90–W97. 10.1093/nar/gkw377

3. Raudvere, U., Kolberg, L., Kuzmin, I., Arak, T., Adler, P., Peterson, H., & Vilo, J. (2019). g:Profiler: a web server for functional enrichment analysis and conversions of gene lists (2019 update). Nucleic Acids Research, 47(W1), W191–W198. 10.1093/nar/gkz369

4. Huang, D.W., Sherman, B.T., Tan, Q., Kir, J., Liu, D., Bryant, D., Guo, Y., Stephens, R., Baseler, M.W., Lane, H.C., & Lempicki, R.A. (2007). DAVID Bioinformatics Resources: expanded annotation database and novel algorithms to better extract biology from large gene lists. Nucleic Acids Research, 35(Web Server issue), W169–W175. 10.1093/nar/gkm415

5. Gene Ontology Consortium. (2023). The Gene Ontology knowledgebase in 2023. Genetics, 224(1), iyad031. 10.1093/genetics/iyad031

6. Evangelista, J.E., Clarke, D.J.B., Xie, Z., Lachmann, A., Jeon, M., Chen, K., Jagodnik, K.M., Jenkins, S.L., Kuleshov, M.V., Wojciechowicz, M.L., Schürer, S.C., Medvedovic, M., & Ma’ayan, A. (2022). SigCom LINCS: data and metadata search engine for a million gene expression signatures. Nucleic Acids Research, 50(W1), W697–W709. 10.1093/nar/gkac328

7. Marino, G.B., Evangelista, J.E., Clarke, D.J.B., & Ma’ayan, A. (2025). L2S2: chemical perturbation and CRISPR KO LINCS L1000 signature search engine. Nucleic Acids Research, 53(W1), W338–W350. 10.1093/nar/gkaf373

8. GTEx Consortium. (2013). The Genotype-Tissue Expression (GTEx) project. Nature Genetics, 45(6), 580–585. 10.1038/ng.2653

9. GTEx Consortium. (2020). The GTEx Consortium atlas of genetic regulatory effects across human tissues. Science, 369(6507), 1318–1330. 10.1126/science.aaz1776

10. Sanford, J.A., Nogiec, C.D., Lindholm, M.E., Adkins, J.N., Amar, D., Dasari, S., Drugan, J.K., Fernández, F.M., Radom-Aizik, S., Schenk, S., Snyder, M.P., Tracy, R.P., Vanderboom, P., Trappe, S., Walsh, M.J., & Molecular Transducers of Physical Activity Consortium. (2020). Molecular Transducers of Physical Activity Consortium (MoTrPAC): Mapping the Dynamic Responses to Exercise. Cell, 181(7), 1464–1474. 10.1016/j.cell.2020.06.004

11. MoTrPAC Study Group. (2024). Temporal dynamics of the multi-omic response to endurance exercise training. Nature, 629, 174–183. 10.1038/s41586-023-06877-w

12. Supek, F., Bošnjak, M., Škunca, N., & Šmuc, T. (2011). REVIGO summarizes and visualizes long lists of gene ontology terms. PLoS ONE, 6(7), e21800. 10.1371/journal.pone.0021800

13. Merico, D., Isserlin, R., Stueker, O., Emili, A., & Bader, G.D. (2010). Enrichment map: a network-based method for gene-set enrichment visualization and interpretation. PLoS ONE, 5(11), e13984. 10.1371/journal.pone.0013984

14. Yu, G., Wang, L.G., Han, Y., & He, Q.Y. (2012). clusterProfiler: an R package for comparing biological themes among gene clusters. OMICS, 16(5), 284–287. 10.1089/omi.2011.0118

15. Yue, Z., Welner, R.S., Willey, C.D., Amin, R., & Chen, J.Y. (2025). GOALS: Gene Ontology Analysis with Layered Shells for Enhanced Functional Insight and Visualization. bioRxiv. 10.1101/2025.04.22.650095

16. Al Abir, F. & Chen, J.Y. (2024). Mondrian Abstraction and Language Model Embeddings for Differential Pathway Analysis. In Proceedings of IEEE International Conference on Bioinformatics and Biomedicine (BIBM), pp. 407–410. IEEE, 2024. 10.1109/BIBM62325.2024.10822686

17. Marino, G.B., Olaiya, S., Evangelista, J.E., Clarke, D.J.B., & Ma’ayan, A. (2025). GeneSetCart: assembling, augmenting, combining, visualizing, and analyzing gene sets. GigaScience, 14, giaf025. 10.1093/gigascience/giaf025

18. Vogelstein, B., Lane, D., & Levine, A.J. (2000). Surfing the p53 network. Nature, 408(6810), 307–310. 10.1038/35042675

19. Simanshu, D.K., Nissley, D.V., & McCormick, F. (2017). RAS proteins and their regulators in human disease. Cell, 170(1), 17–33. 10.1016/j.cell.2017.06.009

20. López-Otín, C., Blasco, M.A., Partridge, L., Serrano, M., & Kroemer, G. (2013). The hallmarks of aging. Cell, 153(6), 1194–1217. 10.1016/j.cell.2013.05.039

21. López-Otín, C., Blasco, M.A., Partridge, L., Serrano, M., & Kroemer, G. (2023). Hallmarks of aging: an expanding universe. Cell, 186(2), 243–278. 10.1016/j.cell.2022.11.001

22. Subramanian, A., Tamayo, P., Mootha, V.K., Mukherjee, S., Ebert, B.L., Gillette, M.A., Paulovich, A., Pomeroy, S.L., Golub, T.R., Lander, E.S., & Mesirov, J.P. (2005). Gene set enrichment analysis: a knowledge-based approach for interpreting genome-wide expression profiles. PNAS, 102(43), 15545–15550. 10.1073/pnas.0506580102

23. Huan, T., Wu, X., & Chen, J.Y. (2010). Systems biology visualization tools for drug target discovery. Expert Opinion on Drug Discovery, 5(5), 425–439. 10.1517/17460441003725102

24. Choi, S. (Ed.). (2010). Systems Biology for Signaling Networks. Springer. 10.1007/978-1-4419-5797-9

25. Huang, H., Wu, X., Pandey, R., Li, J., Zhao, G., Ibrahim, S., & Chen, J.Y. (2012). C2Maps: a network pharmacology database with comprehensive disease-gene-drug connectivity relationships. BMC Genomics, 13(Suppl 6), S17. 10.1186/1471-2164-13-S6-S17

26. Xie, Z., Kropiwnicki, E., Wojciechowicz, M.L., Jagodnik, K.M., Shu, I., Bailey, A., Clarke, D.J.B., Jeon, M., Evangelista, J.E., Kuleshov, M.V., Lachmann, A., Parigi, A.A., Sanchez, J.M., Jenkins, S.L., & Ma’ayan, A. (2022). Getting started with LINCS datasets and tools. Current Protocols, 2, e487. 10.1002/cpz1.487

27. Kanehisa, M., & Goto, S. (2000). KEGG: Kyoto Encyclopedia of Genes and Genomes. Nucleic Acids Research, 28(1), 27–30. 10.1093/nar/28.1.27

28. Gillespie, M., Jassal, B., Stephan, R., Milacic, M., Rothfels, K., Senff-Ribeiro, A., Griss, J., Sevilla, C., Matthews, L., Gong, C., Deng, C., Varusai, T., Ragueneau, E., Haider, Y., May, B., Shamovsky, V., Weiser, J., Brunson, T., Sanati, N., Beckman, L., Shao, X., Fabregat, A., Sidiropoulos, K., Murillo, J., Viteri, G., Cook, J., Shorser, S., Bader, G., Demir, E., Sander, C., Haw, R., Wu, G., Stein, L., Hermjakob, H., & D’Eustachio, P. (2022). The Reactome pathway knowledgebase 2022. Nucleic Acids Research, 50(D1), D986–D992. 10.1093/nar/gkab1028

29. Liberzon, A., Birger, C., Thorvaldsdóttir, H., Ghandi, M., Mesirov, J.P., & Tamayo, P. (2015). The Molecular Signatures Database Hallmark Gene Set Collection. Cell Systems, 1(6), 417–425. 10.1016/j.cels.2015.12.004

30. Miao, Y., Guo, Y., Ma, H., Yan, J., Jiang, F., Liao, R., & Huang, J. (2025). GoBERT: Gene Ontology Graph Informed BERT for Universal Gene Function Prediction. Proceedings of the AAAI Conference on Artificial Intelligence, 39(1), 622–630. 10.1609/aaai.v39i1.32043

31. McInnes, L., Healy, J., & Melville, J. (2018). UMAP: Uniform Manifold Approximation and Projection for Dimension Reduction. arXiv:1802.03426. 10.48550/arXiv.1802.03426

32. Shneiderman, B. (1992). Tree visualization with tree-maps: 2-d space-filling approach. ACM Transactions on Graphics, 11(1), 92–99. 10.1145/102377.115768

