## supplements for "MondrianMap: Navigating Gene Set Hierarchies with Multi-Resolution Enrichment Maps"

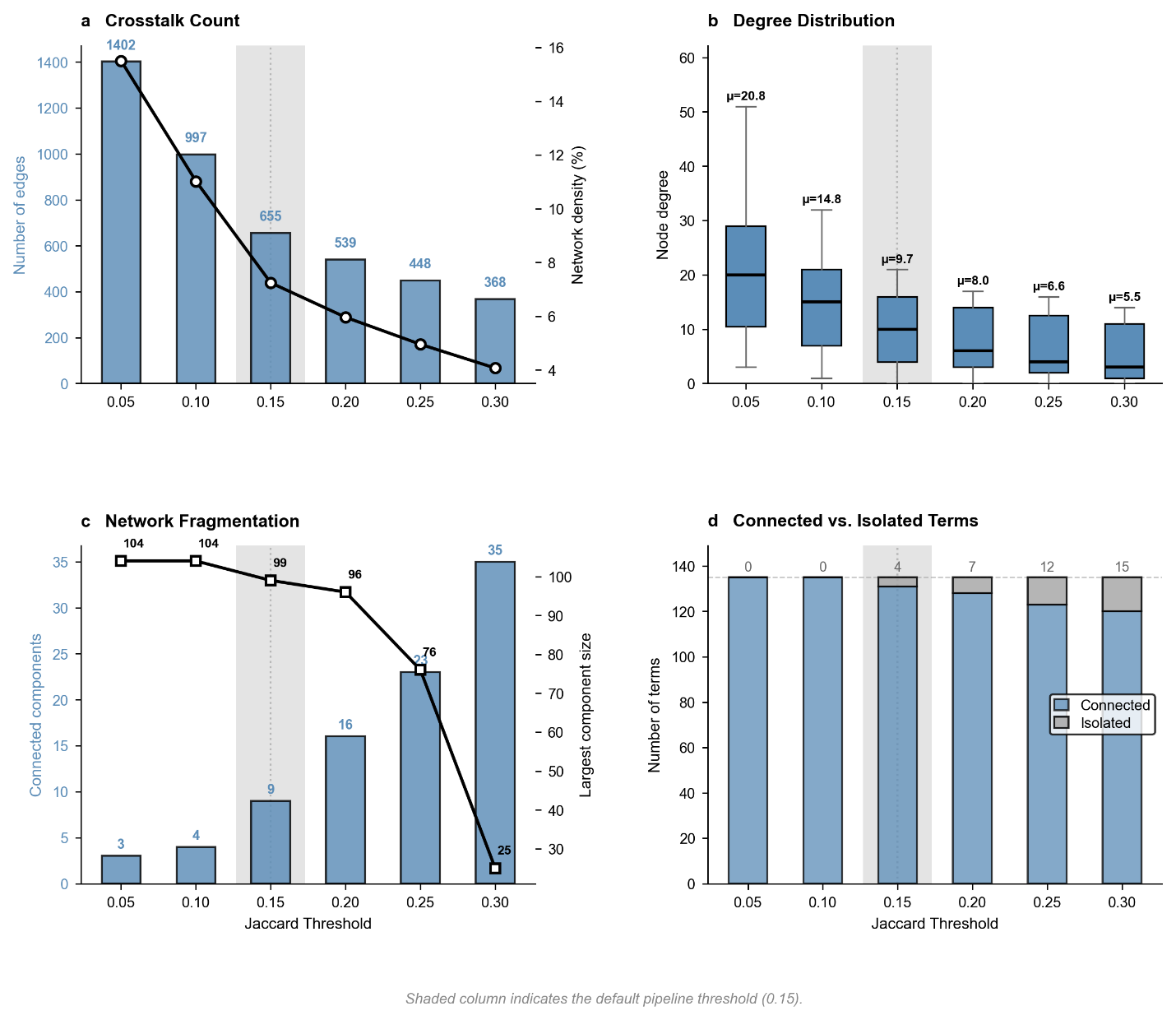

**Figure S1. Jaccard threshold sensitivity for crosstalk network topology.** Network characteristics of gene-set crosstalk edges computed across six Jaccard thresholds (0.05–0.30) for the TP53 CRISPR knockout enrichment in A549 cells K12, 135 GO-BP terms). (a) Total edge count (blue bars, left axis) and network density (orange line, right axis) as a function of the minimum Jaccard index required for an edge. (b) Distribution of node degree at each threshold, with box plots showing median, interquartile range, and outliers; mean degree annotated above each box. (c) Number of connected components (purple bars, left axis) and size of the largest connected component (red line, right axis). (d) Stacked bar chart showing the number of connected versus isolated (degree-zero) terms at each threshold. The shaded column indicates the default pipeline threshold (0.15) used throughout the manuscript.

MondrianMap connects enriched GO-BP terms with crosstalk edges when the Jaccard similarity of their underlying gene sets meets a user-adjustable threshold. Raising the threshold from 0.05 to 0.30 on the TP53 CRISPR knockout enrichment reduced the number of edges from 1,402 to 368 and lowered network density from 15.5% to 4.1% (Figure S1a). Mean node degree dropped from 20.8 at threshold 0.05 to 5.5 at 0.30 (Figure S1b). The default threshold (0.15) occupies a transition zone in the fragmentation curve (Figure S1c–d): it retains 655 edges, keeps 97% of terms connected (only 4 isolates), and yields a mean degree of 9.7. This balance preserves the global connectivity needed for UMAP to embed semantically related terms nearby, while pruning spurious low-overlap edges that would conflate functionally distinct processes.

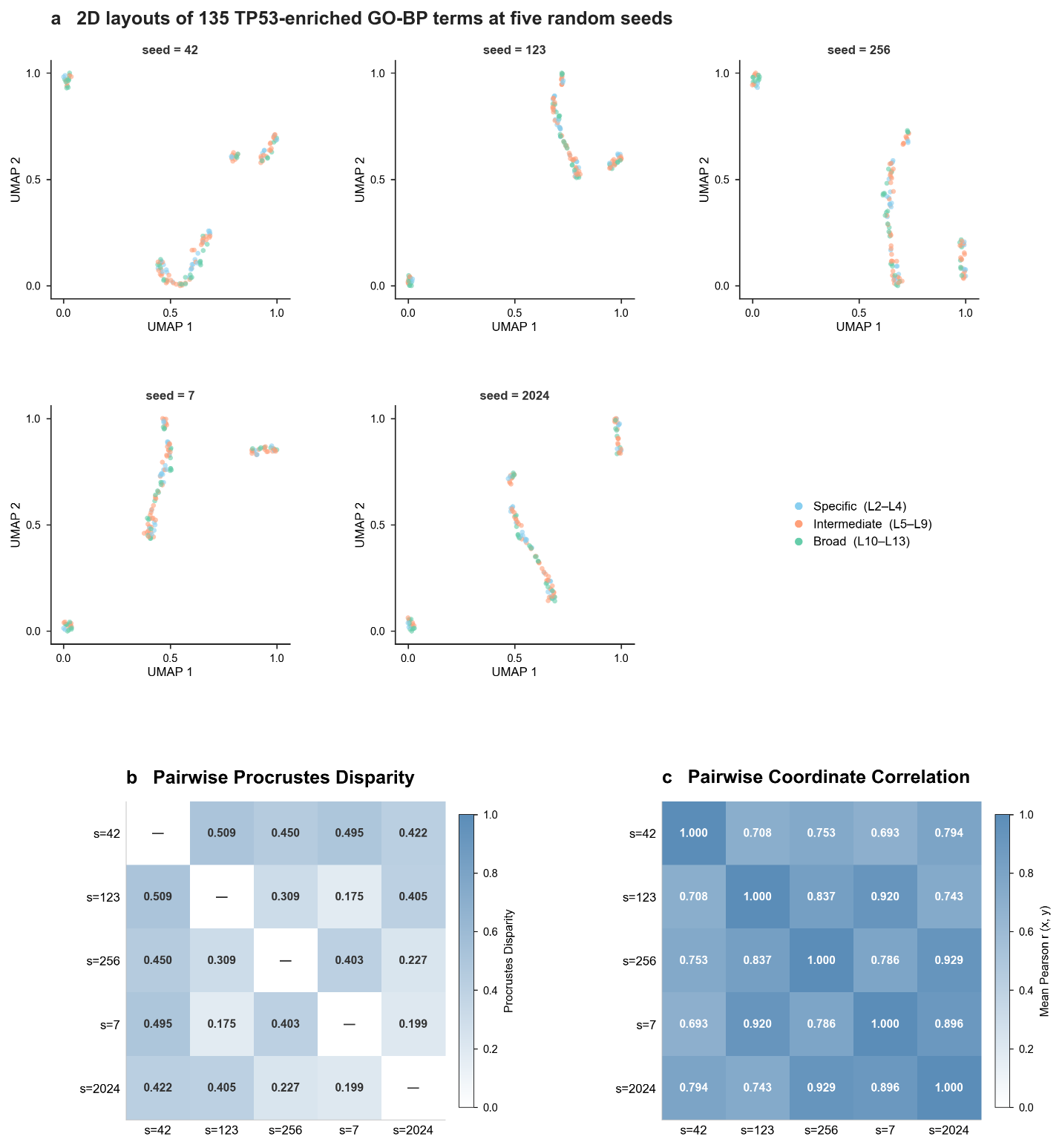

**Figure S2. UMAP layout stability across random seeds.** **(a)** Two-dimensional UMAP projections of 135 TP53-enriched GO-BP terms (GoBERT 1,024-dimensional embeddings, cosine metric) at five random seeds. Scattered points are colored by GOALS tier: Specific (L2–L4, blue), Intermediate (L5–L9, orange), Broad (L10–L13, green). **(b)** Pairwise Procrustes disparity matrix (lower values indicate greater geometric similarity after optimal rotation, translation, and uniform scaling). **(c)** Pairwise Pearson correlation of aligned x- and y-coordinates (mean of the two axes). The summary statistics panel reports aggregate measures across all 10 unique seed pairs.

UMAP is an inherently stochastic algorithm, sensitive to random seed initialization for small sample sizes. To quantify this variability, we projected the GoBERT embeddings of 135 TP53-enriched GO-BP terms into two dimensions at five random seeds (Figure S2). After Procrustes alignment, the mean pairwise disparity was 0.391 ± 0.202, and coordinate correlation averaged r = 0.751 ± 0.152. These results are consistent with the known behavior of UMAP at sample sizes below the n ≈ 500 threshold, where stochastic convergence stabilizes. Crucially, visual inspection (Figure S2a) reveals that the variability manifests primarily as global rotation and local permutation of tightly clustered terms. The coarse-grained cluster structure, the segregation of Specific, Intermediate, and Broad-tier terms, is consistently reproduced across seeds. This confirms that while global orientation may shift, the semantic groupings necessary for accurate biological interpretation remain robust.

**Table S1. TP53 Replicate Concordance for LINCS CRISPR Perturbations**

TP53 knockout (CRISPR) in the A549 cell line at 96 hours post-perturbation. Layer-wise comparison of two independent replicates: K12 (n = 135 enriched GO-BP terms) versus M16 (n = 121 terms). All metrics are computed at the GOALS semantic layers L2–L13.

| Layer | # Terms R1 (K12) | # Terms R2 (M16) | Shared | Term Jaccard | Direction Agreement | Gene Jaccard | Spear ρ | p-value |
| --- | --- | --- | --- | --- | --- | --- | --- | --- |
| L2 | 2 | 1 | 1 | 0.500 | **1.00** | 0.500 | n.d. | n.d. |
| L3 | 9 | 4 | 1 | 0.083 | **1.00** | 0.250 | n.d. | n.d. |
| L4 | 19 | 8 | 5 | 0.227 | **1.00** | 0.122 | 0.368 | 0.542 |
| L5 | 15 | 12 | 3 | 0.125 | **1.00** | 0.203 | 1.000 | 0.000 |
| L6 | 16 | 12 | 4 | 0.167 | **1.00** | 0.294 | 0.800 | 0.200 |
| L7 | 13 | 21 | 8 | 0.308 | 0.75 | 0.109 | 0.455 | 0.257 |
| L8 | 13 | 12 | 4 | 0.190 | **1.00** | 0.235 | 0.949 | 0.051 |
| L9 | 6 | 7 | 3 | 0.300 | **1.00** | 0.268 | 1.000 | 0.000 |
| L10 | 10 | 9 | 4 | 0.267 | **1.00** | 0.203 | –0.800 | 0.200 |
| L11 | 7 | 7 | 2 | 0.167 | **1.00** | 0.122 | n.d. | n.d. |
| L12 | 5 | 3 | 1 | 0.143 | 0.00 | 0.000 | n.d. | n.d. |
| L13 | 20 | 25 | 10 | 0.286 | 0.40 | 0.137 | 0.115 | 0.751 |

**Description**

**# Terms R1 (K12), Terms R2 (M16)**: number of enriched GO-BP terms per replicate at each GOALS layer.

**Shared**: number of GO-BP term identifiers enriched in both replicates at the same layer.

**Term Jaccard**: |shared terms| ÷ |union of terms|, measuring overlap of enriched GO identifiers.

**Direction Agreement**: fraction of shared terms with concordant effect direction (upregulated or downregulated in both replicates).

**Gene Jaccard**: Jaccard similarity of all genes constituting enriched terms per replicate per layer.

**Spear ρ**: Spearman rank correlation coefficient of −log₁₀(adjusted p-value) for shared terms. Computed only when ≥3 shared terms are present at the layer.

**p-value:** two-tailed significance of the Spearman correlation. Reported to 3 significant figures; n.d. = not determined (insufficient shared terms).

**Summary:** Mean Term Jaccard 0.230 ± 0.107; Mean Gene Jaccard 0.204 ± 0.120; Mean Direction Agreement 0.846 ± 0.308. Layers populated: 12 (both replicates). Total enriched terms: 135 (R1) + 121 (R2) = 256.

**Table S2. CS1 LINCS L1000 CRISPR Perturbation Contrasts**

| **#** | **Gene** | **Cell Line** | **Total Terms** | **Up** | **Down** | **Crosstalks** |
| --- | --- | --- | --- | --- | --- | --- |
| 1a | TP53 (R1) | A549 | 135 | 26 | 109 | 655 |
| 1b | TP53 (R2) | A549 | 121 | 11 | 110 | 528 |
| 1c | KRAS (R1) | A549 | 50 | 16 | 34 | 159 |
| 1d | KRAS (R2) | A549 | 33 | 33 | 0 | 72 |
| 1e | MYC | A549 | 27 | 9 | 17 | 76 |
| 1f | EGFR | A549 | 29 | 1 | 28 | 52 |

**Table S3. CS2 GTEx Aging Signatures Contrasts**

| **#** | **Tissue** | **Young Age** | **Old Age** | **Total** | **Up** | **Down** | **Crosstalks** |
| --- | --- | --- | --- | --- | --- | --- | --- |
| 2a | Brain | 20-29 | 60-69 | 63 | 10 | 53 | 223 |
| 2b | Brain | 20-29 | 70-79 | 41 | 13 | 28 | 111 |
| 2c | Muscle | 20-29 | 60-69 | 9 | 0 | 9 | 28 |
| 2d | Muscle | 20-29 | 70-79 | 0 | - | - | - |
| 2e | Heart | 20-29 | 60-69 | 13 | 3 | 10 | 26 |
| 2f | Heart | 20-29 | 70-79 | 22 | 4 | 18 | 52 |
| 2g | Liver | 20-29 | 60-69 | 99 | 0 | 99 | 845 |
| 2h | Blood | 20-29 | 60-69 | 71 | 71 | 0 | 378 |

**Table S4. CS3 MoTrPAC Endurance Trained Rats Contrasts**

| **#** | **Tissue** | **Timepoint** | **Total** | **Up** | **Down** | **Crosstalks** |
| --- | --- | --- | --- | --- | --- | --- |
| 3a | BAT | 1W | 0 | - | - | - |
| 3b | BAT | 2W | 4 | 0 | 4 | 3 |
| 3c | BAT | 4W | 0 | - | - | - |
| 3d | BAT | 8W | 285 | 195 | 90 | 1316 |
| 3e | Heart | 1W | 43 | 43 | 0 | 172 |
| 3f | Heart | 2W | 40 | 1 | 39 | 381 |
| 3g | Heart | 4W | 110 | 0 | 110 | 1108 |
| 3h | Heart | 8W | 0 | - | - | - |
| 3i | Gastroc. | 1W | 36 | 13 | 23 | 121 |
| 3j | Gastroc. | 2W | 24 | 0 | 24 | 157 |
| 3k | Gastroc. | 4W | 13 | 13 | 0 | 24 |
| 3l | Gastroc. | 8W | 19 | 15 | 4 | 43 |
| 3m | Blood | 1W | 26 | 0 | 26 | 94 |
| 3n | Blood | 2W | 4 | 3 | 1 | 3 |
| 3o | Blood | 4W | 0 | - | - | - |
| 3p | Blood | 8W | 42 | 42 | 0 | 81 |
